# Multi-modal cryo-EM reveals trimers of protein A10 to form the palisade layer in poxvirus cores

**DOI:** 10.1101/2023.05.24.541142

**Authors:** Julia Datler, Jesse M Hansen, Andreas Thader, Alois Schlögl, Victor-Valentin Hodirnau, Florian KM Schur

**Affiliations:** Institute of Science and Technology Austria (ISTA), Klosterneuburg, Austria

**Keywords:** Poxvirus, Vaccinia virus, core, cryo-electron microscopy, cryo-electron tomography, AlphaFold

## Abstract

Poxviruses are among the largest double-stranded DNA viruses with members such as Variola virus, Monkeypox virus and the famous vaccination strain Vaccinia virus (VACV). Knowledge about the structural proteins that form the viral core, found in all infectious poxvirus forms, has remained sparse. While major core proteins have been annotated *via* indirect experimental evidence, their structures have remained elusive and they could not be assigned to the individual architectural features of the core. Hence, which proteins constitute which layers of the core, such as the so-called palisade layer and the inner core wall has remained enigmatic.

Here, we have performed a multi-modal cryo-electron microscopy (cryo-EM) approach to elucidate the structural determinants of the VACV core. In combination with molecular modeling using AlphaFold, we unambiguously identify trimers formed by the cleavage product of A10 as the key component of the palisade layer. This allows us to place previously-obtained descriptions of protein interactions within the core wall into perspective and to provide a substantially revised model of poxvirus core architecture. Importantly, we show that interactions within A10 trimers are likely identical among *Poxviridae*, implying that our structural observations should be generalizable over most, if not all members of this important virus family.

**One sentence summary:** Single-particle cryo-EM, cryo-electron tomography, and AlphaFold modeling reveal the structural architecture of the poxvirus core and identify trimers of protein A10 as the key component of the palisade layer.

## Introduction

Poxviruses are large pleomorphic double-stranded DNA viruses infecting a wide range of hosts, from vertebrates, including humans, to arthropods (Condit et al. 2006). Among the members of the *Poxviridae* family are Variola virus, the causative agent of smallpox, and Vaccinia virus (VACV), the prototypical and most extensively studied poxvirus. VACV was also used as an attenuated vaccination strain to eradicate smallpox in the late 1970s (Strassburg 1982). Recently, the re-emergence of Monkeypox virus causing localized outbreaks of Mpox around the globe re-emphasized the importance to gain a better understanding of the intricate poxvirus lifecycle.

Poxvirus replication occurs within viral factories that are exclusively located within the cytoplasm of a host cell and gives rise to immature viruses (IVs), which then eventually transition into different forms of infectious mature virions (MVs) (Roberts and Smith 2008). MVs are enveloped by a lipid bilayer and contain a dumbbell-shaped core encapsulating the viral DNA genome, and so-called lateral bodies (LB) which laterally attach to the exterior of the core wall and contain viral proteins for modulating host immunity and oxidative response (Condit et al. 2006, Bidgood and Mercer 2015, Bidgood et al. 2022) (**Figure 1A**).The transition from IVs to MVs requires the proteolytic cleavage of several core proteins which in turn contribute to the formation and condensation of the viral core, with its characteristic shape and biochemical signature (Ansarah-Sobrinho and Moss 2004, Liu et al. 2014). As the core is one of the uniting factors in all the infectious poxvirus forms and fulfills one of the key roles in the virus lifecycle, i.e. the protected transfer of the viral genome and required accessory proteins to a newly infected cell, substantial effort has been invested into its detailed structural and biochemical characterization. However, the structural determinants that underlie core morphogenesis have remained poorly understood, impeded by the molecular complexity of poxviruses and the apparent lack of sequence homology of the suggested structural protein candidates to other species.

**Figure 1:**
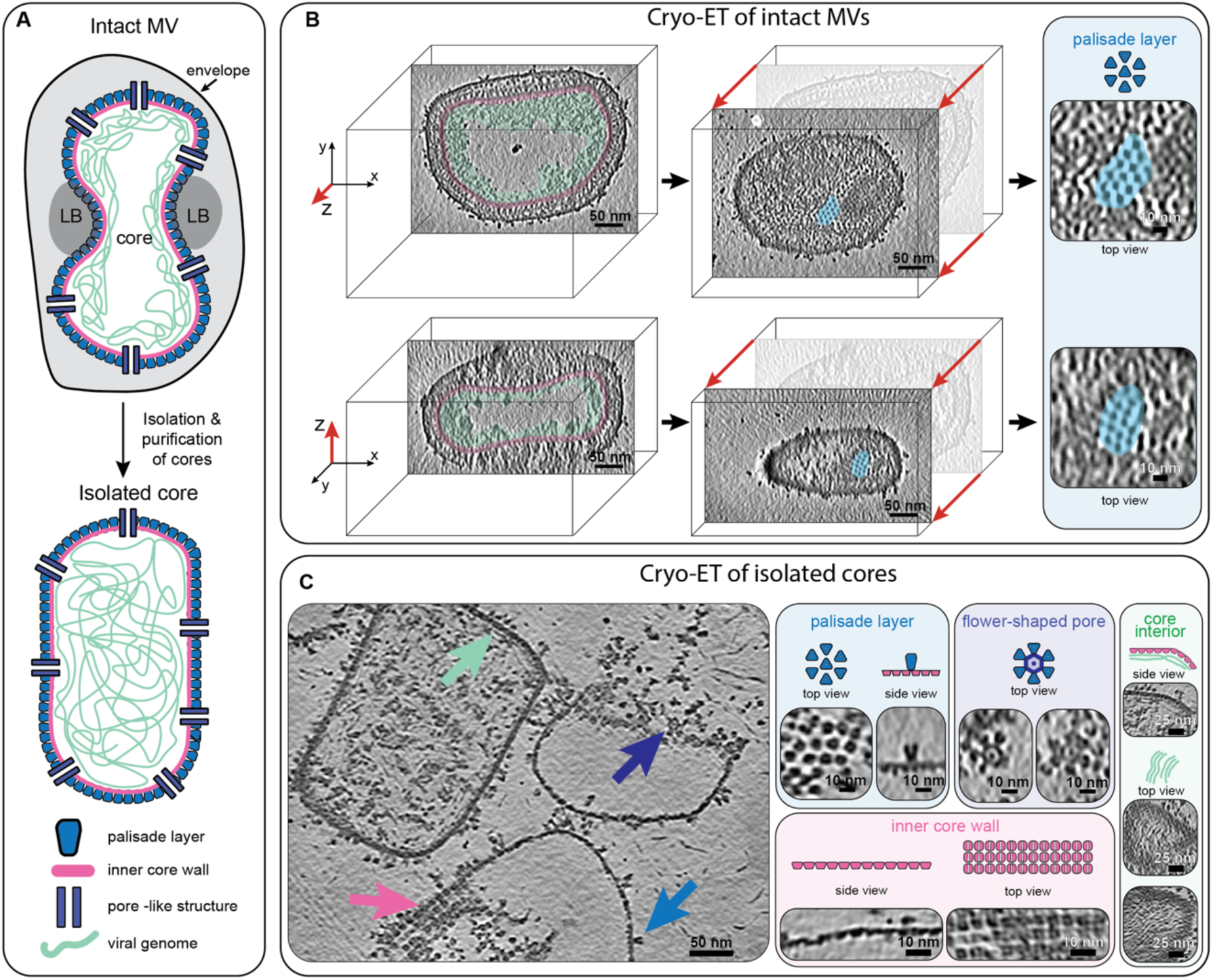
Cryo-ET of in VACV mature virions and isolated cores. **A)** Schematic depiction of intact MV (top) and an isolated MV core (bottom), showing previously described structural entities, such as palisade layer, inner core wall, pore-like structure and viral genome. Lateral bodies (LB) bind to the concave-shaped core in intact viruses, but are often lost during core isolation. This color coding for the palisade layer (blue), the inner core wall (pink), the pore-like structure (purple) and the genome (green) is kept consistent throughout subsequent figures. **B)** Computational slices (1.1nm thickness) through a missing wedge-corrected tomogram using IsoNet (Liu et al. 2022) of an intact MV particle clearly showing core morphology and structural features, such as the hexameric arrangement of the palisade layer (highlighted in blue), the inner core wall (pink) and the condensed genome (green). The virus is shown from two viewing directions looking at the xy and xz planes (see axes on left) on top and bottom, respectively. The center panel shows the palisade layer in grazing slices, with a magnified view of this region shown on the right. **C)** Computational slice (1.1nm thickness) through an IsoNet-corrected tomogram of isolated MV cores (left). Structural features are clearly observable in tomograms of isolated cores (annotated with arrows, same color scheme as A) and are shown on the right. Scale bar dimensions are annotated in the figure.

Room-temperature and cryo-electron microscopy (cryo-EM) analysis of VACV revealed the presence of a regular, but discontinuous, palisade layer formed of spike-like assemblies on the outside of a continuous viral inner core wall (Dubochet et al. 1994, Pedersen et al. 2000, Moussatche and Condit 2015). Improved imaging modalities using cryo-electron tomography (cryo-ET) further suggested the spikes of the palisade layer to be forming a pseudo-hexameric lattice (Cyrklaff et al. 2005, Hernandez-Gonzalez et al. 2023). In addition, several studies suggested the presence of pore-like structures with unknown function spanning the core wall (Cyrklaff et al. 2005, Moussatche and Condit 2015, Hernandez-Gonzalez et al. 2023).

Beyond these morphological descriptions of the core architecture, identities of proteins forming the core wall and palisade layer were derived from studies using 1) immunogold labeling of intact VACV MV and purified cores (Vanslyke and Hruby 1994, Cudmore et al. 1996, Roos et al. 1996, Risco et al. 1999, Pedersen et al. 2000, Moussatche and Condit 2015), 2) biochemical and proteomic studies via extraction and partitioning of MV components (Jensen et al. 1996, Chung et al. 2006, Mirzakhanyan and Gershon 2019, Bidgood et al. 2022) and 3) genetic experiments via recombinant viruses and inducible mutants of selected protein candidates (Risco et al. 1999, Heljasvaara et al. 2001, Rodriguez et al. 2006, Jesus et al. 2014, 2015). Specifically, protocols to isolate intact cores from native virus particles (**Figure 1A**), using non-ionic detergent under reducing conditions, allowed a more straightforward biochemical and ultrastructural description (Dubochet et al. 1994, Moussatche and Condit 2015, Bidgood et al. 2022).

Together these studies provided an inventory of the proteins presumably forming the core wall and palisade layer, namely the cleavage products of precursor protein A10 (p4a in the UniProt database), precursor protein A3 (p4b in the UniProt database), A4 (p39 in the UniProt database) and L4 (VP8 in the UniProt database) (**Figure S1A,B**). For consistency with current use of protein nomenclature in the poxvirus field we will exclusively refer to the core proteins as A10, A3, A4 and L4. The minor cleavage product of A10 will be referred to as 23K.

In immunogold labeling experiments A10 and A4 were detected at the outer surface of the core wall (Cudmore et al. 1996, Roos et al. 1996, Risco et al. 1999, Pedersen et al. 2000, Moussatche and Condit 2015). Proteins A3 and A4 were named as components of the palisade layer (Wilton et al. 1995, Cudmore et al. 1996). Peculiarly, protein A3 was also suggested to be located at the inner part of the core wall (Moussatche and Condit 2015). Furthermore, immunolabelling of cryosections (Pedersen et al. 2000) and broken viral cores (Moussatche and Condit 2015) located L4 at the inner core wall, in line with its role as a major DNA binding protein (Bayliss and Smith 1997, Jesus et al. 2014).

However, given these also partially contradictory results, no direct structural proof could be obtained to unambiguously assign any of these protein candidates to specific structural core features, i.e. the palisade layer, the inner core wall or protein densities in the core interior. Without this knowledge, key steps of the poxvirus lifecycle remain enigmatic, limiting possibilities for extending the potential tools for pharmacological interference during poxvirus infection.

Here we have used a combination of cryo-ET, subtomogram averaging (STA) and single particle cryo-electron microscopy (cryo-SPA) to study complete VACV MVs and isolated VACV virus cores. Our results show that the palisade layer and inner core wall adopt two different local symmetries and we identify several novel distinct structural entities in the core. Importantly, our integrated use of cryo-ET and SPA, combined with AlphaFold (Jumper et al. 2021) identifies trimers of A10 to form the palisade layer. Together, these results allow us to extend the structural atlas of poxvirus cores and present a substantially refined model of the poxvirus core assembly.

## Results

### Cryo-ET of VACV mature virus

No experimentally-derived structures of the major structural core protein candidates are available. Hence, we used AlphaFold (Jumper et al. 2021) to computationally predict models of the main core protein candidates, A10, 23K, A3, A4, and L4 (**Figure S1C**) to facilitate the interpretation of cryo-EM densities obtained in our downstream workflow.

In a first attempt to structurally annotate the proteins forming the individual features of the viral core we performed high-resolution cryo-ET on intact VACV MV virions purified from infected HeLa cells (**Table S1**). Despite the relatively large virus dimensions (∼360nm x 250nm x 220nm), our reconstructed tomograms allowed us to visualize fine details of virus particles with their characteristic brick-shaped overall morphology and dumbbell-shaped core (**Figure 1B, Movie S1**). In line with previous cryo-EM studies of MVs (Cyrklaff et al. 2005, Hernandez-Gonzalez et al. 2023), the exterior of the core surface was coated with spikes of the palisade layer. The core lumen was predominantly empty, except for the condensed viral genome underlying the inner core wall. In order to obtain high-resolution structures of the individual layers of the core we performed STA (**Figure S2A**). This revealed the palisade layer to be composed of hexamer of trimers (**Figure S2B**), as suggested previously (Hernandez-Gonzalez et al. 2023). In our STA structure, we did not observe clearly ordered densities for the inner core wall, suggesting that it is not following the same arrangement as the palisade layer. At this point our structures obtained from intact virus particles had insufficient resolution (∼13Å) (**Figure S2C**) to identify the proteins forming the trimers or the inner core wall. However, our alignment protocol still allowed us to clearly visualize the overall arrangement of the palisade layer into a large-scale pseudo-hexagonal lattice (**Figure S2D**). The lattice displayed large areas of continuous organization interspersed with gaps and cracks breaking the lattice into locally symmetric patches. We did not observe any obvious pentamer formation which could allow the complete closure of a hexagonal lattice. This is reminiscent of the incomplete hexagonal lattice observed, for example, with Gag proteins in immature retroviruses (Mattei et al. 2016). Using the initial mesh defined on the surface of the core wall for extracting subtomograms, and the measured size of a trimer within a hexamer of trimer unit, we calculated an average number of ∼2280 trimers to constitute the palisade layer, not considering the presence of gaps and cracks.

### Cryo-ET of isolated MV cores reveals their structural complexity

In order to improve the resolution of core structural features, we decided to reduce the complexity of our experimental system and therefore isolated VACV cores via optimizing established protocols (Joklik 1962, Esteban 1984, Dubochet et al. 1994). Dubochet and colleagues showed that trimers can be released from isolated cores and visualized as isolated particles upon vitrification (Dubochet et al. 1994). In our vitrified sample, isolated cores sometimes partially collapsed and appeared seemingly empty, as reported previously (Dubochet et al. 1994). However, usually they retained a regular barrel shape, not displaying the concavities observed within MVs (**Figure 1C, Movie S2**). The genome filled the entirety of the viral core (**Figure 1C, left core**) rather than being restricted to just the region underneath the inner core wall as seen in intact MVs (**Figure 1B**), indicating that core isolation potentially leads to genome decondensation.

Strikingly, in partially collapsed cores a multitude of structural features was visible already in the individual tomograms, which we could, given the 3D nature of tomograms, assign to the individual layers of the core (**Figure 1C**). In particular, we could visualize trimers of the palisade layer, seemingly still organized into a pseudo-hexagonal lattice. More importantly, we could for the first time observe the structural arrangement of the inner core wall. Each inner core wall unit adopted a square-like shape of ∼7.4nm x 7.4nm and appeared to be following at least 2-fold symmetry, which is consistent with our interpretation from intact viruses that the inner core wall is not following the organization of the palisade layer. As the third main architectural feature, we found flower-shaped structures (diameter ∼29nm), consisting of flower ‘petals’ and a central ring with hexameric symmetry and an inner diameter of ∼11nm. Our cryo-ET data suggested the outer petals of these flower-like assemblies to each be comprised of a palisade trimer. Positioned within the hexameric center we consistently observed a strong density. These assemblies may be the core wall pores reported in previous lower-resolution cryo-ET data (Cyrklaff et al. 2005, Hernandez-Gonzalez et al. 2023) and negative stain EM images of isolated cores (Moussatche and Condit 2015). Directly below the inner core wall linear densities could often be observed, organized into a parallel striated pattern (**Figure 1C**). The dimensions of these densities suggest them to be DNA.

### SPA reveals a diversity of structures in isolated cores

Given the high-quality of the isolated cores and clear visibility of structural features, we reasoned that SPA cryo-EM would allow us to further improve the resolution of our structures. Hence, we acquired two SPA datasets (**Figure 2A, B**). One still contained isolated viral cores and the second only retained individual components released from cores as the sample was prepared with an additional centrifugation purification step prior to vitrification. 2D classification of this data revealed the structural treasure chest of the VACV core (**Figure 2C**), yielding classes of trimeric, tetrameric, pentameric, and hexameric assemblies. Beyond classes for soluble particles, released either from the core wall or the core interior, we obtained classes of the flower-shaped pores which were still retained in the core wall (**Figure 2A**). In addition, we obtained classes for continuous segments of the core wall. Some classes contained only the inner core wall, resembling in appearance the two-fold symmetric assembly observed in tomograms of isolated cores. Other classes contained the inner core wall plus densities of the trimers in the palisade layer. Notably, we could not obtain classes where both the inner core wall and the palisade layer were both equally well resolved, again indicating an independent organization of these two structural layers with respect to each other. Given the wealth of structures within our sample, we set out to determine their higher-resolution structures.

**Figure 2:**
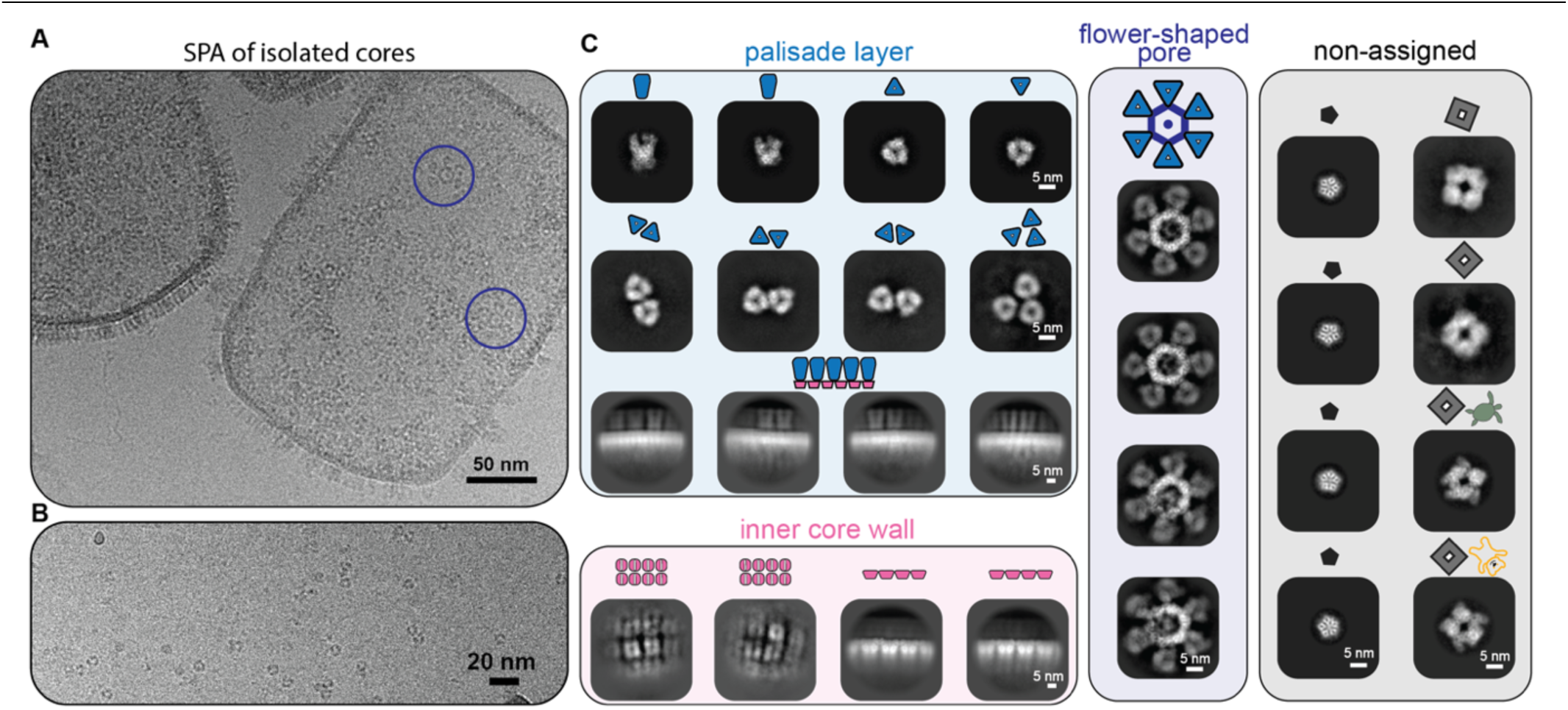
The structural treasure chest of isolated VACV cores. **A-B)** Example micrographs of the SPA acquisition showing isolated cores (A) and soluble core proteins (B), such as trimers. The shown micrographs are from the same data acquisition. The flower-shaped pore also observed in cryo-ET of isolated cores (Figure 1C) is highlighted in purple circles. **C)** Gallery of 2D classes obtained from processing cryo-SPA data sets showing different multimeric assemblies observed as part of the core or as soluble components. Comparison of these 2D classes with our cryo-ET data allows a clear contextualization of their origin with respect to the palisade layer, the inner core wall, or the flower-shaped pore. Classes for which no features could be observed in our cryo-ET data are labeled as non-assigned. The assembly and symmetry state of the classes is also annotated with small schematic depictions. Scale bar dimensions are annotated in the figure.

### Proteomics of isolated cores

First, to verify that our sample preparation retained protein candidates of interest we performed mass spectrometry of the soluble protein fraction in the isolated core sample (**Table S2**). Our proteomics data confirmed that our SPA sample was enriched for the major structural core proteins A10, A3, A4 and L4. In addition, we found many other proteins previously reported to be packaged into the core, including proteins involved in transcription and translation of the viral genome as well as several host proteins.

### Trimers of A10 constitute the palisade layer

For structure determination, we first focused on the trimers in our SPA data, given their prominent structural appearance as part of the palisade layer. Using 3D refinement in RELION, we obtained a high-resolution reconstruction of the trimer at a global resolution of 4.2Å, which we further improved using Phenix’s density modification (Terwilliger et al. 2020) to a global resolution of 3.8Å (**Figure 3A-B, S3**). This map allowed the visualization of structural detail such as bulky side chains and alpha-helical pitch (**Figure 3B**) and enabled precise fitting of our computationally predicted models (**Figure S1C**) into our EM density. This unambiguously revealed three copies of protein A10 (modeled residues 1-599) to form the trimer (**Figure 3C, Movie S3**).

**Figure 3.**
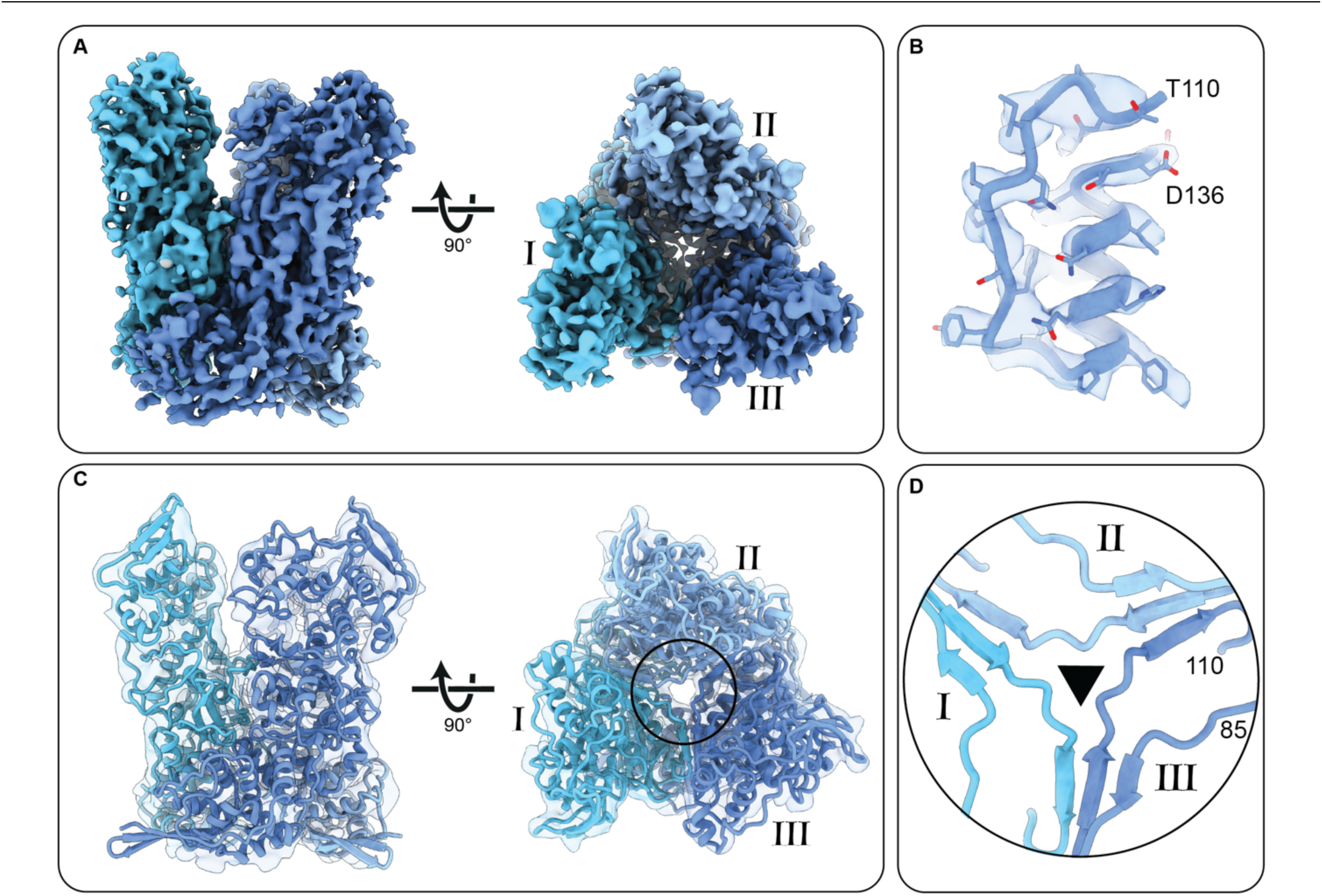
Single particle cryo-EM structure of A10 trimer. **A)** C3-symmetric density-modified cryo-EM reconstruction of A10 trimer at 3.8Å resolution. Each subunit of the trimer (I to III) is depicted with a different shade of blue. **B)** Highlight of a core region in the A10 trimer cryo-EM map where side chain density permits verification of primary protein sequence. **C)** Refined model fit into cryo-EM density for the A10 trimer. The circle shows the zoomed-in region of interest in panel D. The transparent EM-density has been low-pass filtered to 10Å resolution to facilitate interpretation. **D)** Central trimer contacts from panel C showing residues 85-110 which engage in hetero-oligomer beta-sheet interactions with neighboring monomers. The residue numbers of the N-terminus and C-terminus of the displayed protein region are annotated.

### The A10 trimer is stabilized via an extensive oligomerization interaction network

Next, we used the PDBePISA server (Krissinel and Henrick 2007) to calculate properties and identify putative key contacts at the A10 trimer interface (**Figure S4, MovieS4**). We note that at ∼4Å resolution structural features, such as small and/or negatively charged side chains are not clearly visible and hence impose a limit to interpretability. This analysis revealed that hydrophobic interactions dominate within the 2123Å^2^ buried surface area per protomer pair (**Figure S4A**). There are also several inter-chain salt bridges tethering central alpha helices together within the trimer (**Figure S4B**). Remarkably, each pair of protomers assembles a core heterodimeric three-stranded beta-sheet (residues 85-110, **Figure 3C, Movie S4**) formed by two strands from one monomer and one strand from the neighboring monomer. A hydrogen bond network on the outward surface of the beta-sheet further reinforces this interaction, while the opposite side of the beta-sheet packs tightly with underlying hydrophobic side chains (**Figure S4C**). Finally, our analysis revealed a plausible disulfide bridge between residues C32 and C569 (**Figure S4D**) which conceivably clamps the upper and lower halves of one A10 monomer together.

### The palisade layer forms only weak lateral interactions between trimers

The model of our trimer fits into our STA density map with high agreement, leaving no major area of density within the palisade layer unoccupied (**Figure S5A**). This strongly suggests that the A10 trimer is the main constituent of the palisade layer. However, our fit into the STA map also revealed that the lateral interactions across trimers within the palisade layer are not extensive given the wide spacing between them. This further implies that the stabilization of the core is not entirely contributed by the A10 trimer but probably depends on the underlying inner core wall and additional interactions above the trimer.

When further examining our lattice maps of the palisade layer obtained via STA, we observed a variable orientation of the individual trimers with respect to each other (**Figure S5B**). This agrees with the existence of variable multimeric trimer classes in the soluble fraction of the isolated core SPA data set (**Figure 2C, Figure S5B**), where we observed interacting trimers with substantial differences in their positioning with respect to each other. Together, these two findings suggest that the variable inter-trimer orientation within our lattice maps, which fits with the weak lateral palisade layer interactions we describe, likely represents a finding with biological significance.

### Structural positioning of other core wall proteins with respect to the trimer

Given that A10 is forming most, if not all of the palisade layer, we wondered how other structural proteins of the core wall may interact with our A10 trimer structure so that we could develop a spatial model of wall organization. We mapped previously reported mass spectrometry cross-linkages (Mirzakhanyan and Gershon 2019) onto our trimer model (**Figure S6A**). A4 appears to interact preferentially with the exterior side of the A10 trimer, in line with previous immunogold labeling experiments (Pedersen et al. 2000, Moussatche and Condit 2015). 23K and L4 interactions map throughout the trimer model, though L4 tends to link more with centrally-located residues. The major core protein A3 forms linkages exclusively with the interior side of the A10 trimer, suggesting that A3 could be a component of the inner core wall density we observe in our tomograms and SPA 2D class averages. Interestingly, the bottom of the A10 trimer facing the inner core wall is strongly positively charged (**FigureS6B**). Reciprocally, A3 has a negatively charged patch at one of its sides (**FigureS6C**). Indeed, when we generated a low-resolution structure of the inner core wall density from our SPA data, an AlphaFold prediction of an A3 dimer fit into this density with good agreement (**Figure S7**).

### The flower-shaped pore of the core wall

Next, we focused on the hexameric flower-shaped pore, which was positioned at the same height as the trimer within the core layer, but also further extended above as described (Hernandez-Gonzalez et al. 2023) (**Figure 1C,2C**). Our cryo-EM reconstruction at ∼7Å (**Figure S8, S9**) revealed the outer densities surrounding the center indeed to be the trimers of the palisade layer (**Figure S9A**). Due to map anisotropy we were unable to unambiguously fit a model into the hexameric density at the core. However, in both C6 and C1 reconstructions, we again observed an unidentified donut-shaped density on the outward-facing side of the pore (**Figure S9C**). The fact that interaction between trimers and the hexameric center of the pore is not affected by core isolation and does not lead to shedding of trimers within the petals suggests that these interactions must be stronger than those between adjacent trimers in the palisade layer.

### The A10 trimer is conserved across poxvirus species

The protein sequence of A10 is highly conserved among the orthopoxvirus genus, including the key residues forming interactions in the trimer, with an average ∼97% sequence identity between VACV Western reserve (WR), Variola virus, Monkeypox virus, Rabbitpox virus, Cowpox virus and Ectromelia virus (**Figure S10A**). Correspondingly, AlphaFold predictions of these proteins as monomers or trimers are very similar (**Figure S10B**). This strongly suggests that the interactions formed within the trimers constituting the palisade layer are identical among *Poxviridae* and imply that our structural observations should be generalizable over most, if not all members of this virus family.

## Discussion

By using a combination of SPA to obtain a high-resolution structure of the trimer forming the palisade layer, and cryo-ET to contextualize our observations of novel structural entities within VACV cores, we have substantially extended the understanding of poxvirus core architecture.

We have unambiguously identified A10 as the key protein forming the palisade layer of VACV cores, which now provides the possibility to place previously obtained descriptions of protein interactions and locations within the core wall into perspective and to provide a revised model of poxvirus core architecture (**Figure 4**).

**Figure 4:**
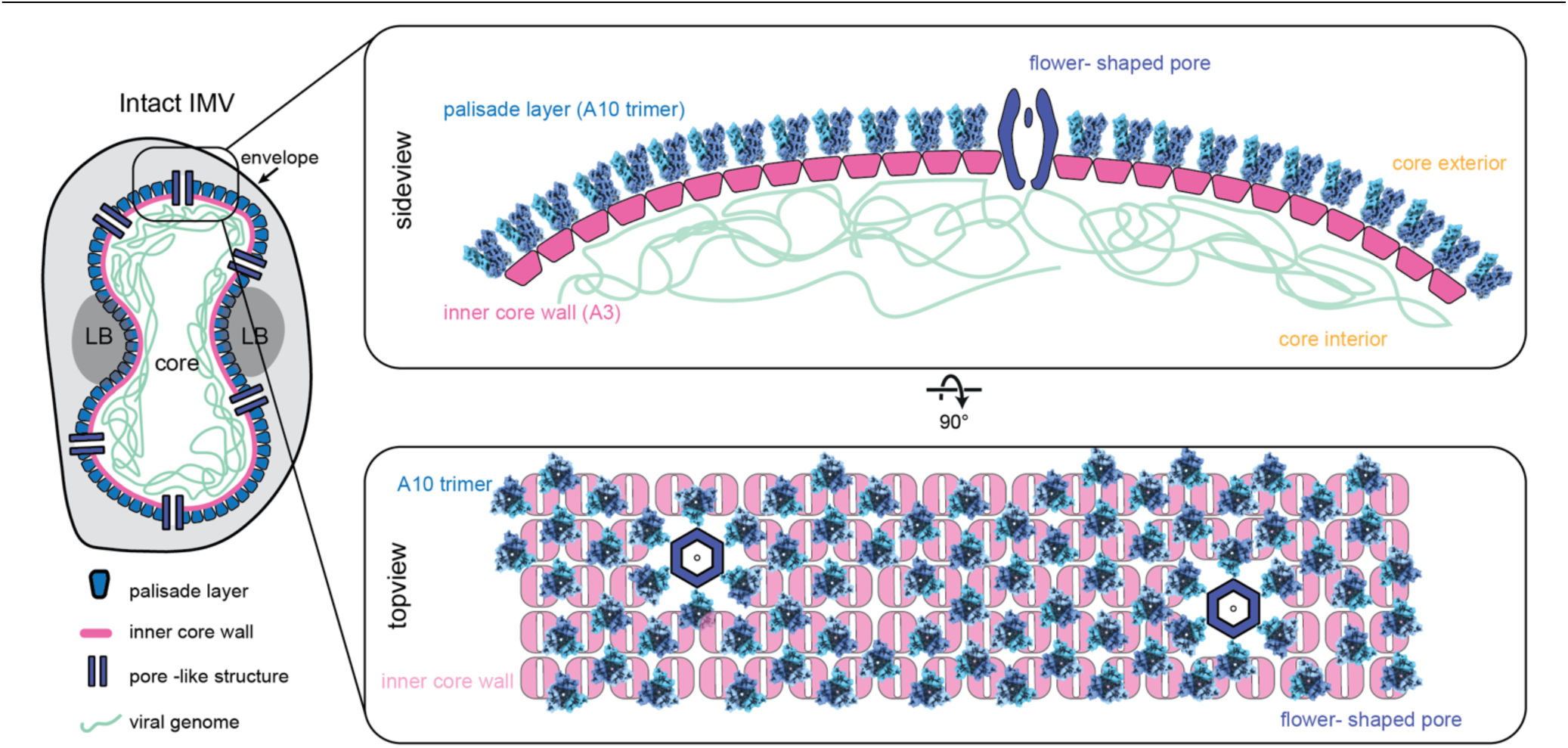
Structural model of the VACV core wall. Schematic summary of the updated model of the palisade layer and inner core wall. Protein A10 forms the palisade layer, which is positioned above an inner core wall with strikingly different symmetry. A4 is most likely decorating the outside of the palisade layer. The inner core wall is presumably formed by A3, with a potential role of L4 as a DNA-binding protein, tethering the viral genome to the core wall. The core is pierced by flower – shaped pores, which appear unevenly distributed on the surface of the core.

### The position of structural proteins within the core

Immunoprecipitation experiments showed A10 and A4 to form a stable complex even before proteolytic cleavage and MV formation (Risco et al. 1999). Moreover, previous studies using immunolabeling speculated the spike protein of the palisade layer to be A4 (Cudmore et al. 1996, Roos et al. 1996, Pedersen et al. 2000). We show that A4 is not part of the trimers in the palisade layer, which is in line with its disordered nature revealed by AlphaFold prediction **(Figure S1C)**. Instead, interaction sites between A4 and A10 are located preferentially on the core exterior-facing side of the trimer (Mirzakhanyan and Gershon 2019) (**Figure S6A**). This is consistent with a report claiming A4 to be on the exterior of the core based on data showing A4 to be partially disappearing after purification of the cores with the detergent NP40 and DTT (Roos et al. 1996, Moussatche and Condit 2015). Hence, we consider it likely that A4, while providing some stabilizing role to the palisade layer, could rather be acting as a matrix-like protein, for example establishing a link between the core and the surrounding membranes as described previously (Cudmore et al. 1996), or to be part of the pore.

Cross-linking mass spectrometry (XL-MS) data of VACV cores also suggested a direct interaction between protein A10 and A3 (Mirzakhanyan and Gershon 2019). Based on our model, these linkages are exclusively positioned on the bottom side of the trimer facing the core interior, suggesting that A3 is part of the inner core wall **(Figure S6A).** This is again in line with previous experiments which revealed that A3 is only detectable via immunogold labeling upon breakage of the core after hypertonic shock and protease treatment (Moussatche and Condit 2015). Accordingly, our low-resolution map of the inner core wall accommodates a dimer of A3. A small region of unoccupied density on the top of the inner core wall facing the palisade layer (**Figure S7D, dashed rectangle**), might be occupied by 23K, given its placement in the proximity of the trimer based on XLMS data. It is tempting to speculate about potential interaction interfaces between A10, 23K and A3 proteins based on this observation, but more experimental work is needed to unambiguously define the positioning and the interactions between the individual core wall proteins. L4 is a major DNA binding protein (Bayliss and Smith 1997, Jesus et al. 2014) and has been reported to be located at the inside of the viral core (Pedersen et al. 2000, Moussatche and Condit 2015). In this regard, the predicted interactions between L4 and A10 based on XL-MS (Mirzakhanyan and Gershon 2019) **(Figure S6A)** are not conclusive.

Still, taking these observations into account, we deem the most likely arrangement of the major core proteins positions to be A4 on the outside of the viral core, A10 forming the palisade layer, as revealed in this study, and A3 building the inner core wall. Given that there is no unoccupied density in the palisade layer, 23K could be positioned either below the trimer or in the pore. Whether L4 might be also located in the pore hexagon or its central density remains to be identified.

### A potential role of A10 as a shape-defining structural protein in the core wall

The finding of A10 forming trimers that constitute the palisade layer raises the question of their exact function. Our observation that the lateral interactions between trimers are not extensive and result in variable inter-trimer interactions within the lattice, argues against a pure core-stabilizing role. The integrity of the viral core must therefore be achieved by another layer, such as the inner core wall, or aided by A4 that could be linking the trimers on the exterior of the viral core (Cudmore et al. 1996). Instead, it is tempting to speculate that the trimers in the palisade layer are acting as shape determinators of the core, allowing it to form the highly complex dumbbell-shaped structure observed in MV, with both convex and concave curvatures. In line with this hypothesis are conditional mutation studies that showed that upon loss of A10, the correct assembly to MVs is not possible (Rodriguez et al. 2006) and instead, the inner core wall builds stacks or sheet-like architectures labeled by L4, F17 and E8 antibodies (Heljasvaara et al. 2001). This may suggest that A10 provides curvature-defining attributes to the viral core, while the inner core wall could act as a stabilizer via stronger lateral protein-protein interactions. This is also reflected in isolated cores, where trimers can be shed off the inner core wall, while the core itself is held intact by the inner core wall. A similar shape-defining role of a viral protein can be found in retroviral capsid proteins, where its subdomains have been proposed to be either providing curvature-defining features or to be the main stabilizing protein-protein interactions (Schur et al. 2015).

### The poxvirus core pore

The flower-shaped pore identified in our cryo-EM and cryo-ET datasets most likely corresponds to the previously described pore-like structure (Cyrklaff et al. 2005, Moussatche and Condit 2015, Hernandez-Gonzalez et al. 2023). Its function was postulated to be either directly involved in the mRNA extrusion into the cytoplasm of infected cells (Cyrklaff et al. 2005, Moussatche and Condit 2015) or to be the hexameric rings of the viral D5 primase/helicase, which is essential for Vaccinia virus genome release (Hernandez-Gonzalez et al. 2023). Our cryo-EM density shows an additional density in the center of the flower lumen. Theoretically, the lumen would be large enough to accommodate DNA, but further experimental proof will be required to show if the pore plays a role in transporting nucleic acids through the core wall.

### Structural characterization of novel protein structures from virus lysates

By working with isolated cores, which effectively should contain all the components the virus packs within it for infecting new cells, we were able to visualize several structural entities with varying symmetries. Given their abundance, they most likely play a relevant role in the viral lifecycle (**Table S1**). This includes the pentamer, which was highly abundant in our data, and the tetrameric structures (**Figure 2C**).

Despite considerable efforts, we were not able to resolve structures, aside from the trimers, to high enough resolutions to unambiguously assign their identities. Future work could change the purification protocols of cores, and improve sample vitrification protocols to reduce preferred orientation, in order to obtain higher-resolution structures of the other components.

*In vitro* reconstitution of structural core proteins could also be pursued for the unidentified core classes. For example, this has been successfully used for structural studies of D13 which forms a hexameric lattice of the immature virus (Bahar et al. 2011, Hyun et al. 2011, 2022). *In vitro*, reconstitution of A10 trimers or other oligomers would be an interesting avenue to further elucidate the complex structural assembly mechanism of the mature poxvirus core.

In a concurrent preprint by Liu & Corroyer-Dulmot et al., the authors also show A10 to form trimers within the palisade layer. Classification of trimers on the surface of isolated cores further reveals an increased flexibility of the trimer, in line with our observations of variable inter-trimer interactions. Given the recent re-emergence of the Monkeypox virus causing global epidemics, our and the concurrent work provide important fundamental new insights into the poxvirus core architecture.

## Acknowledgments

We thank Andreas Bergthaler (Research Center for Molecular Medicine of the Austrian Academy of Sciences) for providing VACV WR. We thank Armel Nicholas and his team at the ISTA proteomics facility, and Stefano Elefante at the ISTA Scientific Computing facility for their support. We also thank Florian Fäßler, Darío Porley, Theresa Muthspiel and other members of the Schur group for support and helpful discussions. We also thank Daniel Castaño-Díez for support with Dynamo.

FKMS acknowledges support from ISTA and EMBO. FKMS also acknowledges the support from the Chan Zuckerberg Initiative. This research was also supported by the Scientific Service Units (SSUs) of ISTA through resources provided by Scientific Computing (SciComp), the Life Science Facility (LSF), and the Electron Microscopy Facility (EMF).We also acknowledge the use of COSMIC (Cianfrocco et al. 2017) and Colabfold (Mirdita et al. 2022).

## Author contributions

Project administration: F.K.M.S.; supervision and funding acquisition: F.K.M.S.; conceptualization: J.D. and F.K.M.S.; methodology: J.D., J.M.H. and F.K.M.S.; investigation: J.D., J.M.H., A.T., V.-V.H., A.S.; Software: A.S; validation, formal analysis, and visualization: J.D., J.M.H. and F.K.M.S.; data curation: J.D., J.M.H. and F.K.M.S.; writing—original draft: J.D., J.M.H. and F.K.M.S.; writing—review and editing: J.D., J.M.H., A.T., V.-V.H., A.S and F.K.M.S

## Data availability

The electron microscopy density maps of the A10 trimer and the hexameric flower-shaped pore, the subtomogram average of the palisade layer, as well as representative tomograms for complete viruses as well as isolated cores have been deposited in the Electron Microscopy Data Bank under accession codes: EMD-17410, EMD-17411, EMD-17412, EMD-17413, EMD-17414. The refined model of the A10 trimer has been deposited in the Protein Data Bank accession code: PDB 8P4K.

## Material and Methods

### Virus propagation and purification

Vaccinia virus Western Reserve (VACV WR) was kindly received from Andreas Bergthaler (CeMM & Medical University of Vienna). VACV WR stock was trypsinized in a 1:1 dilution with 0.25% Trypsin (Thermo Fisher Scientific, no. 25200056) for 30 min at 37°C and 5% CO2. HeLa cells (kindly received from the Sixt lab, ISTA) were seeded in Dulbecco’s modified Eagle medium GlutaMAX (DMEM) (Thermo Fisher Scientific, no. 31966047), supplemented with 10% (v/v) fetal bovine serum (FBS) (Thermo Fisher Scientific, no. 10270106) and 1% (v/v) penicillin-streptomycin (Thermo Fisher Scientific, no. 15070063) in T-175 flasks (Corning, no. CLS431080). Cells were washed once with PBS and infected with seed virus diluted in 2ml infection medium (DMEM, 2.5% FBS, 1% penicillin-streptomycin) at an ∼ multiplicity of infection (MOI) of 1.5 and incubated for 2 hours at 37°C and 5% CO2 with manual shaking every 30 min. After 2 hours, 33ml of infection medium was added and after 3 days, when a cytopathic effect was visible, cells were harvested by scraping. The cell suspension was centrifuged at 1,200xg for 10 min at 4°C and the pellet was resuspended in 500µl 10mM Tris-HCl, pH 9 (Carl Roth, no.9090.3) buffer. The cell pellet containing MV’s was stored at −80°C and cells were lysed by 3x freeze and thaw cycles. Samples were centrifuged for 5 min at 300xg at 4°C and supernatant was collected. The cell pellet was resuspended in 500µl 10mM Tris-HCl buffer pH 9, centrifuged again and supernatant was again collected. Supernatants were pooled and applied on a 6ml sucrose cushion of 36% sucrose (Sigma-Aldrich, no. 84100) in 10mM Tris-HCl buffer, pH 9 in small tubes (Thermo Scientific, Thin-Walled WX, no.03699) and centrifuged at 32,900xg, at 4°C for 80 min in an ultracentrifuge (Sorvall, WX100+, Rotor TH-641). The pellet was resuspended in 500µl 1mM Tris-HCl, pH9 and applied on a sucrose gradient (40%, 36%, 32%, 28%, 24% in 1mM Tris-HCl buffer pH 9) and centrifuged at 26,000xg at 4°C for 50 min (Sorvall, WX100+, Rotor TH-641). The virus, visible via a milky band, was collected and sedimented in a fresh tube with 10ml 1mM Tris-HCl buffer pH 9, at 15,000xg, 30 min at 4°C (Sorvall, WX100+, Rotor TH-641). The purified virus pellet was dissolved in 100µl 1mM Tris-HCl, aliquoted and stored at −80°C until further use.

### VACV core purification

VACV core purification was optimized from previously described protocols (Easterbrook 1966, Dubochet et al. 1994). As described above, the milky band in the sucrose gradient containing the virus was collected and sedimented in a fresh tube with 10ml 1mM Tris-HCl buffer pH 9 at 15,000xg for 30 min at 4°C, and then dissolved in 500µl core stripping buffer containing 0.1% NP-40 (Thermo Scientific, no.85124), 50mM DTT (Carl Roth, no.69083), 50mM Tris-HCl pH 9 and 2U DNAse (Promega, no. M6101). The virus was incubated for 10 min at room temperature and then centrifuged at 20,000xg, 30 min, at 4°C, through a 2ml 24% sucrose cushion in 1mM Tris-HCl, pH 9. Viral cores were collected in 500µl 1mM Tris-HCl buffer, pH 9 and sedimented at 15,000xg for 30 min at 4°C. The final pellet was resuspended in 50µl 1mM Tris-HCl and frozen in aliquots at −80°C until further use. For cryo-SPA samples of isolated VACV WR cores, four times more virus was used during the purification and 3M KCl was added to a final concentration of 300mM before freezing at −80°C.

### Purification of soluble fraction from isolated cores

Aliquots were thawed on ice and 3M KCl was added to a final concentration of 250mM after all following dilution steps. Samples were mixed 1:3 with 0.25% Trypsin and sonicated 3x 30s in a sonication bath (Elma, Elmasonic S40) at 4°C with the sweeping option. After 30 min incubation at 37°C the samples were sonicated again 3 x 30s. Samples were frozen at −80°C and thawed shortly at 37°C and sonicated again at 4°C 3x 30s. Samples for cryo-SPA were centrifuged with 3,488g for 45 min and the supernatant was used for fixation and vitrification as described below. Samples for mass spectrometry were filtered through a 0.1µm filter (Ultrafree MC-VV, Durapore PVDF 0.1um, Merck). The filter was prewetted with 1M Tris-HCl and centrifuged for 3 min, at 12,000xg at 4°C. Flow through was discarded and the sample was added and centrifuged again. Flow through was stored at −80°C until further use.

### Virus cryo-EM preparation and fixation

Whole VACV WR mature viruses and isolated cores were thawed on ice. For cryo-SPA samples of isolated VACV WR cores, 3 M KCl was added after thawing resulting in a final concentration of 210mM KCl after all of the following dilutions steps. All samples except the purified soluble fraction, were sonicated 3x 30s in a sonication bath at 4°C with the sweeping option. 0.25% Trypsin was added in a 1:1 dilution and incubated for 30 min at 37°C. All samples were fixed 1:1 with 4% PFA (Merck, no. P6148) (final concentration 2% PFA) in 1 mM Tris-HCl buffer, pH 9 and incubated 30 min at room temperature and 30 min at 37°C to inactivate the samples. Samples were frozen again at −80°C and then sonicated 3x 30s in a sonication bath at 4°C with the sweeping option before vitrification.

### Cryo-electron microscopy

10nm BSA-Gold (Aurion Immuno Gold Reagents, no. 410.011) in PBS was added to the cryo-ET samples in a dilution of 1:10. Samples for cryo-ET were deposited onto 300-mesh holey carbon grids (Quantifoil Micro Tools, R2/2 X-103-Cu300) and samples for cryo-SPA were deposited onto 200-mesh holey carbon grids (Quantifoil Micro Tools, R 2/2 X-103-Cu200) which were first glow-discharged for 2.5 min using an ELMO glow discharge unit (Cordouan Technologies). 2.5μl of the sample was always added to both sides of the grid, which were vitrified using back-side blotting in a Leica GP2 plunger (Leica Microsystems). Blotting chamber conditions were 80% humidity and 4°C. The grids were vitrified in liquid ethane (−185°C) and then stored under liquid nitrogen conditions until imaging.

Singe particle and tomography data sets were acquired under cryogenic conditions on an FEI Titan Krios G3i TEM microscope (Thermo Fisher Scientific) operating at 300kV and equipped with a Bioquantum post-column energy filter and a Gatan K3 direct detector.

Cryo-electron tomography data was collected with the SerialEM software package version 3.8 (Mastronarde 2005). New gain reference images were collected before data acquisition. DigitalMicrograph 3.4.3 as integrated into the Gatan Microscopy Suite v3.3 (Gatan) was used for filter tuning and SerialEM for microscope tuning. Tilt series were acquired with a filter slit width of 10 eV, using a dose-symmetric tilt scheme (Hagen et al. 2017) ranging from −66° to 66° with a 3° increment. The nominal defocus range was set from −1.5 to −8 μm for the whole VACV WR mature virions and from −1.5 to −5μm for the isolated viral cores. The nominal magnification was set to 64,000×, resulting in a pixel size of 1.381Å. Tilt images were acquired as 5760 × 4092 pixel movies of ten frames. The cumulative dose over the entire tilt series was 165 e/Å^2^. For data acquisition settings, see Table S1.

The automated collection for the isolated viral core cryo-SPA dataset was set up using EPU version 2.13 (Thermo Fisher Scientific) in conjunction with AFIS. The soluble fraction purified from isolated cores was acquired via SerialEM version 4.0 (Mastronarde 2005) with an active beam tilt/astigmatism compensation. SPA micrographs were acquired in counting-mode with a filter slit width of 20 eV, and using a 4-shot per hole data collection. The nominal defocus was set from −1.25 to −3μm for isolated cores and from −1.5 to −2.2μm for the soluble fraction. The nominal magnification was set to 81,000×, resulting in a pixel size of 1.06Å.

The isolated viral core dataset was acquired with 5760 × 4092 pixel movies of 34 frames with a cumulative dose of 53.06 e/Å^2^. The soluble fraction dataset was acquired with a tilted stage of 25 degrees as 5760 × 4092 pixel movies of 54 frames with a cumulative dose of 80.20 e/Å^2^. The decision to acquire tilted data was based on results from the isolated core dataset, which revealed preferential orientation of several of the classes. Details for data acquisition can be found in Table S1.

### Image processing cryo-ET

The image processing workflow is schematically displayed in Figure S2A. Tomoman was used to sort and create stacks (Wan 2020). Defocus was estimated using CTFFIND4 (Rohou and Grigorieff 2015). IMOD (Kremer et al. 1996) was used for tilt series alignment and to generate 8x binned tomograms with weighted back projection. The full tomograms were reconstructed in NovaCTF (Turoňová et al. 2017) with simultaneous 3D CTF correction with a slab thickness of 15nm using the phase flip algorithm.

Bin8 tomograms were then filtered with IsoNet (Liu et al. 2022) to obtain better contrast and also fill missing information in z, in order to allow better visualization of the core **(Figure S11).** Definition of subtomogram averaging starting positions and all subsequent subtomogram averaging steps were performed in Dynamo version 1.1.333 (Castaño-Díez et al. 2012). As starting positions for subtomogram averaging we defined a mesh following the surface of viral cores in IsoNet-corrected bin8 tomograms. To generate a *de novo* reference, subtomograms (cubic size 464 Å³) were then extracted from IsoNet-corrected bin8 tomograms and subjected to five rounds of alignment applying no symmetry.

The obtained reference, which already displayed the hexamer of trimers arrangement described by Hernandez-Gonzalez et al. 2023 was then used to start a new subtomogram alignment in bin8 with subtomograms extracted at the initial mesh positions from weighted back projected tomograms (not IsoNet-corrected). C3 symmetry was applied again only after the 3-fold symmetry of the structure became clearly apparent upon initial bin8 iterations. Alignment was gradually refined from bin8 over bin4 (subtomogram cubic size 464Å³) to bin2 (subtomogram cubic size 398Å³) while advancing the low-pass filter and decreasing the Euler angle scanning step and range. After the first two alignments in bin8, lattices were space cleaned (6 pixels) and cc-threshold cleaned to remove subvolumes that did not align to the core surface. At the stage of bin2, the dataset was split into even/odd halfsets and from this stage on, even/odd datasets were treated absolutely independently. Up to this point the low-pass filter never extended beyond 25Å. After the final bin2 iteration, the final halfset averages were multiplied with a Gaussian-filtered cylindrical mask and the resolution was determined by mask-corrected Fourier-shell correlation (Chen et al. 2013). The final map was sharpened with an empirically determined B-factor of −2100A^2^ and filtered to its measured resolution (Rosenthal and Henderson 2003).

### Image Processing cryo-SPA

Movies from the dataset containing intact cores were motion-corrected with dose-weighting using the RELION 4.0 (Scheres 2012) implementation of MotionCorr2 with a patch size of 7 x 5. Motion-corrected micrographs were then imported into Cryosparc 4.0.0 (Punjani et al. 2017) for subsequent processing. Processing details are summarised here and full details are also presented in Figure S3 (processing of the trimer) and Figure S8 (processing of the flower-shaped pore). Initial CTF parameters were estimated using CryoSPARC patch CTF, and initial picks were obtained using a blob picker. Particles were extracted with a large box size (636Å, bin2) early on to capture both large and small protein populations during 2D classification. Iterative 2D classification with varying mask sizes permitted sorting of particles into different protein species with distinct classes. Particles were then re-extracted with a more appropriate box size: 340Å for trimers, 545Å for the flower-shaped pore, and 636Å for the side views of the core wall. In all cases, initial 3D volumes were generated using CryosSPARC *ab initio* without symmetry. Symmetry was then imposed during 3D auto-refinement.

Selected 2D classes containing particles for the flower-shaped pore were reconstructed using CryoSPARC non-uniform refinement, then locally sharpened and filtered within CryoSPARC. For the 3D reconstruction of the inner core, 3,795 particles containing side views with clearest features of the inner core wall were selected and an initial model was generated using CryoSPARC *ab initio*. The output was used as an initial model for non-uniform 3D refinement without masking and without symmetry imposed. The final map is un-sharpened. Particles of the A10 trimer were exported to RELION for further processing: 2D classification, 3D refinement using a mask containing the full trimer density, Bayesian polishing, defocus refinement, and focused 3D classification of A10 monomers. Importantly, we manually balanced particle views of the trimer by removing over-represented top views during 2D classification, significantly improving map quality. Finally, the resulting map was density modified (Terwilliger et al. 2020) using as input two half maps and the mask used during refinement. 3DFSC calculations were made using the Remote 3DFSC Processing Server (https://3dfsc.salk.edu). Projections of the final flower-shaped pore were made using the V4 tool in EMAN/1.9. (Ludtke et al. 1999).

The dataset collected at 25° stage tilt containing the soluble fraction was motion corrected in RELION (Scheres 2012) identically to the dataset above. Data were processed separately from the dataset containing intact cores. CryoSPARC (Punjani et al. 2017) patch CTF was used to estimate CTF to account for stage tilt. Particles were picked with blob picker as described above. Particles were extracted at bin2 with a box size of 340Å and subjected to iterative 2D classification with varying mask size and 250 classes.

### AlphaFold prediction of core protein candidates

Initial predicted structures of putative structural core proteins, either as monomer or multimers, were generated using Alphafold 2.3.2. (Jumper et al. 2021). Five seeds were generated per model and one prediction per seed. All five models were relaxed using Amber relaxation and inspected manually. The highest-ranking model as determined by pLDDT score was selected for analysis. The following proteins were folded as monomers: A10 (UniProt. P16715, residues 1 - 614), A3 (UniProt. P06440, residues 62 - 643), 23K (UniProt. P16715, residues 698 - 891), A4 (UniProt. P29191) and L4 (UniProt. P03295, residues 33 - 251) **(Figure S1)**. Multimer predictions were done with identical settings to monomer folds. The multimer prediction of the A10 trimer (UniProt. P16715, residues 1 - 614) and the predictions for the comparison of A10 trimer of Variola virus (GeneID ABF23487.1, residues 1 - 615) and Monkeypox virus (GeneID YP_010377118.1, residues 1 - 614) to VACV WR UniProt. P16715, residues 1 - 614) as well as the multimer prediction of A3 dimer (UniProt. P06440, residues 62 - 643) were done also with the AlphaFold version 2.3.2.

### Model building for the A10 trimer

Three copies of the highest-ranking model were rigid-body fit into the density-modified trimer map shown **in Figure 3A**. The A10 trimer was relaxed into the density-modified trimer *via* centroid relaxation in Rosetta/3.13 (20220812 build) (Fleishman et al. 2011, Khatib et al. 2011, Maguire et al. 2021). The model was then imported into Coot (ver. 0.8.9.1 EL) (Emsley et al. 2010) for per-residue manual inspection on a single chain and real-space refinement if necessary. To ensure the symmetry of the final model, this chain was copied to other C3 symmetry positions using Chimera (Pettersen et al. 2004). Model statistics were calculated using the online Molprobity tool (http://molprobity.biochem.duke.edu/) (Williams et al. 2018) with hydrogens first removed and then added back to the model. Pairwise RMSD calculations were done using ChimeraX (Pettersen et al. 2021) matchmaker tool. One final trimer model was rigid-body fit into the outer density of the flower-shaped pore and symmetrized 6-fold using ChimeraX (Pettersen et al. 2021).

### Data visualization and figure preparation spell out

Cryo-electron tomograms of whole VACV WR mature viruses and isolated cores as well as the lattice maps were visualized in IMOD (Kremer et al. 1996), UCSF Chimera (Pettersen et al. 2004) and UCSF ChimeraX (Pettersen et al. 2021). EM-densities were displayed in ChimeraX (Pettersen et al. 2021). Figures were prepared using Adobe Illustrator (Adobe Inc.). Movies were generated in ChimeraX (Pettersen et al. 2021) and Adobe Premiere Pro 2023 (Adobe Inc.).

### Proteomics

#### Sample preparation

To the soluble fraction from isolated cores (prepared as described above) 25mM TCEP (Gold Biotechnology, no. 51805-45-9) and 4% SDS (Carl Roth, no. 8029,1) was added and boiled 10 min at 95°C and was first cleaned up by SP3 using a commercial kit (PreOmics GmbH, 100 mg of beads per sample), then processed using the iST kit (PreOmics GmbH) according to the manufacturer’s instructions. Tryptic digestion was stopped after 1 hour and samples were vacuum dried and then re-dissolved in the iST kit’s LC LOAD buffer with 10 min sonication.

#### LC-MS/MS analysis

The sample was analyzed by LC-MS/MS on an Ultimate 3000 RSLC_Nano nano-HPLC (ThermoFisher Scientific) coupled with a Q Exactive HF (ThermoFisher Scientific), concentrated over an Acclaim PepMap C18 pre-column (5µm particle size, 0.3mm ID x 5 mm length, ThermoFisher Scientific), then bound to an EasySpray C18 column (2 µm particle size, 75 µm ID x 50 cm length, ThermoFisher Scientific) and eluted over the following 60 min gradient: solvent A, MS-grade H₂O + 0.1% formic acid; solvent B, 80% acetonitrile in H₂O + 0.08% formic acid; constant 300 nL/min flow; B percentage: 5 min, 1%; 45 min, 31%; 65 min, 44%. Mass spectra were acquired in positive mode with a Data Independent Acquisition (DIA) method: FWHM 8 s, MS1 parameters: Centroid mode, 1 microscan, 120,000 resolution, AGC target 3e6, 50 ms maximum IT, 400 to 1005 m/z; DIA scans: 24 MS2 scans per cycle, 57 windows of 11.0 m/z width per cycle covering the range from 394.9319 to 1022.21204 m/z (−0.005 m/z non-covered gap between adjacent windows), spectra acquired in Profile mode, with 1 microscan, at 30,000 resolution; AGC target 1e6, 60 ms maximum IT, NCE 27.

#### Data analysis

The raw file was searched in DIANN version 1.8.1 in library-free mode against Homo sapiens and Vaccinia virus (strain Western Reserve) proteomes sourced from UniprotKB. Match-Between-Runs was turned off. Fixed cysteine modification was set to Carbamidomethylation. Variable modifications were set to Oxidation (M) and Acetyl (protein N-term). Data were filtered at 1% FDR.

DIANN’s output was re-processed using in-house R scripts, starting from the report table. Peptide-to-protein assignments were checked, then Protein Groups were assembled and quantified.

---

**Figure S1:**
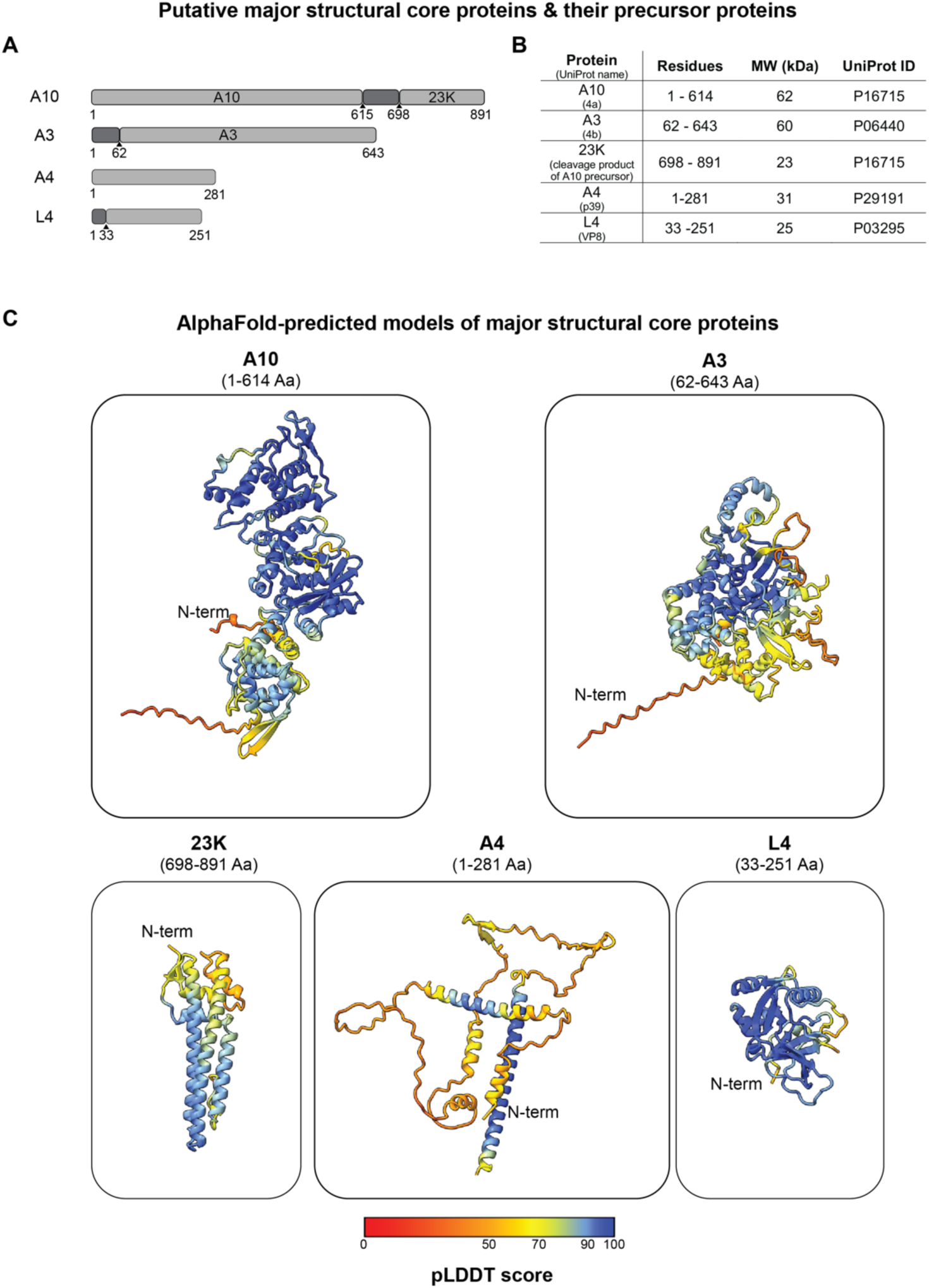
Major structural proteins annotated to be in the VACV core wall. **A)** Schematic representation of VACV WR proteins reported to constitute the core wall and palisade layer. Cleavage sites within precursor proteins are annotated by arrows and the resulting cleavage proteins are listed within the schematic representation. **B)** List of the structural core proteins discussed in this manuscript (in their cleaved form when applicable), with information on their residue length, molecular weight (kiloDalton) and UniProtID. **C)** AlphaFold-predicted (Jumper et al. 2021) models of A10, A3, 23K, A4 and L4. The modeled residues are indicated in brackets. The coloring of the model reflects the pLDDT confidence score (red = low confidence, blue = high confidence).

**Figure S2:**
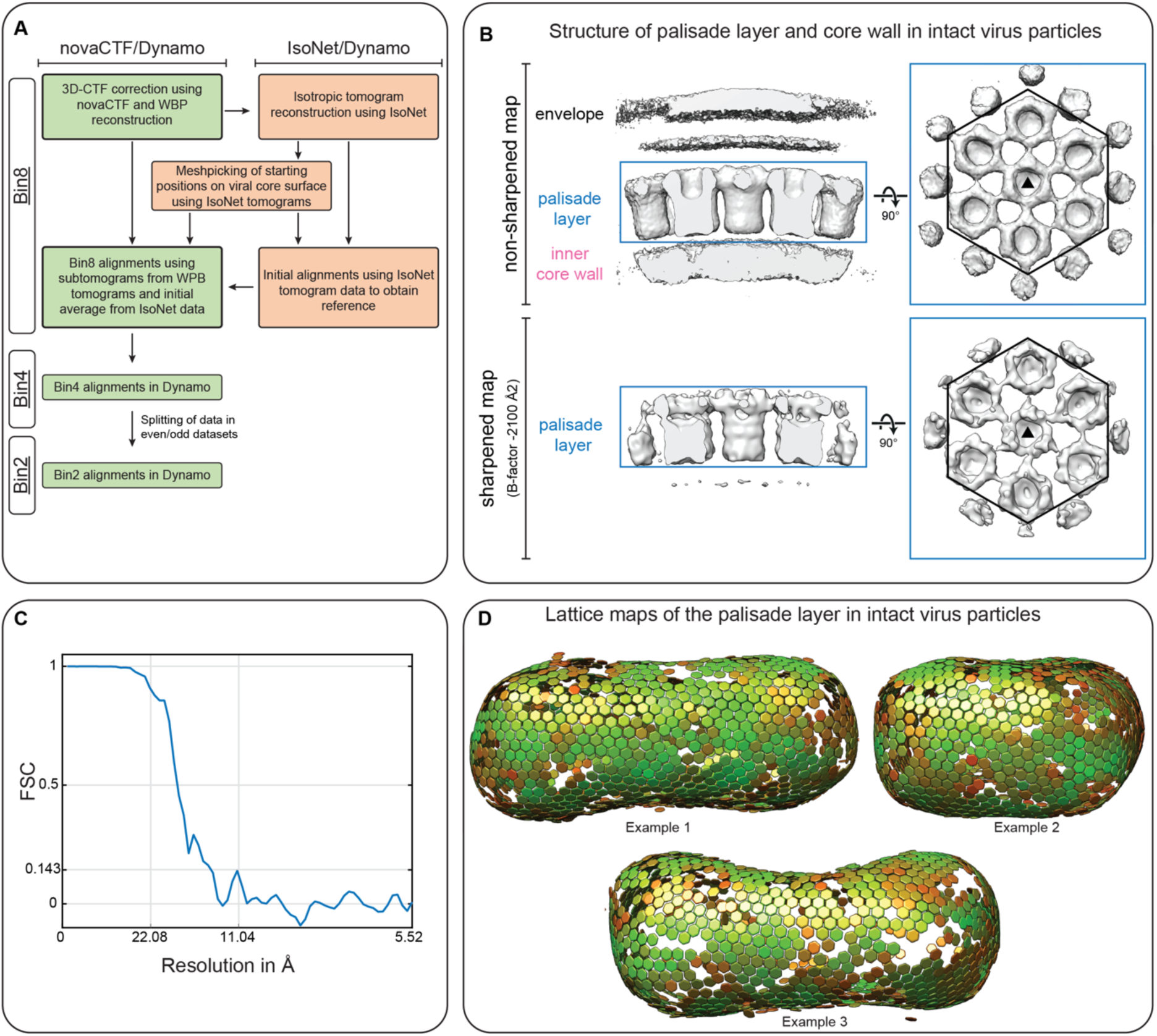
Subtomogram averaging of the VACV core wall in intact MV. **A)** Subtomogram averaging workflow of the core in intact MV using novaCTF (Turoňová et al. 2017), Isonet (Liu et al. 2022) and Dynamo (Castaño-Díez et al. 2012). See Materials and Methods for more details. **B)** Non-sharpened map (top) and sharpened EM-density map (bottom) of the subtomogram average of the palisade layer and inner core wall in intact virus particles. The map is shown in a side view (left) and as seen from outside of the virus (right), where the envelope layer is removed to allow a clear view of the palisade layer. The triangle symbol annotates the central trimer and the hexagon annotates the hexamer of trimers. **C)** Fourier Shell correlation (FSC) between independent half datasets of the map shown in B. The measured resolution at the 0.143 criterion is 13.1Å. **D)** Examples of viral core lattices after final subtomogram averaging alignments. The aligned subtomogram positions are shown as hexagons to allow a facilitated interpretation of the results. Please note that each hexagon position actually represents the trimeric center of a hexamer of trimers. The color of the hexagons denotes the cross-correlation coefficient (CCC) of the alignment ranging from red (low CCC) to green (high CCC).

**Figure S3:**
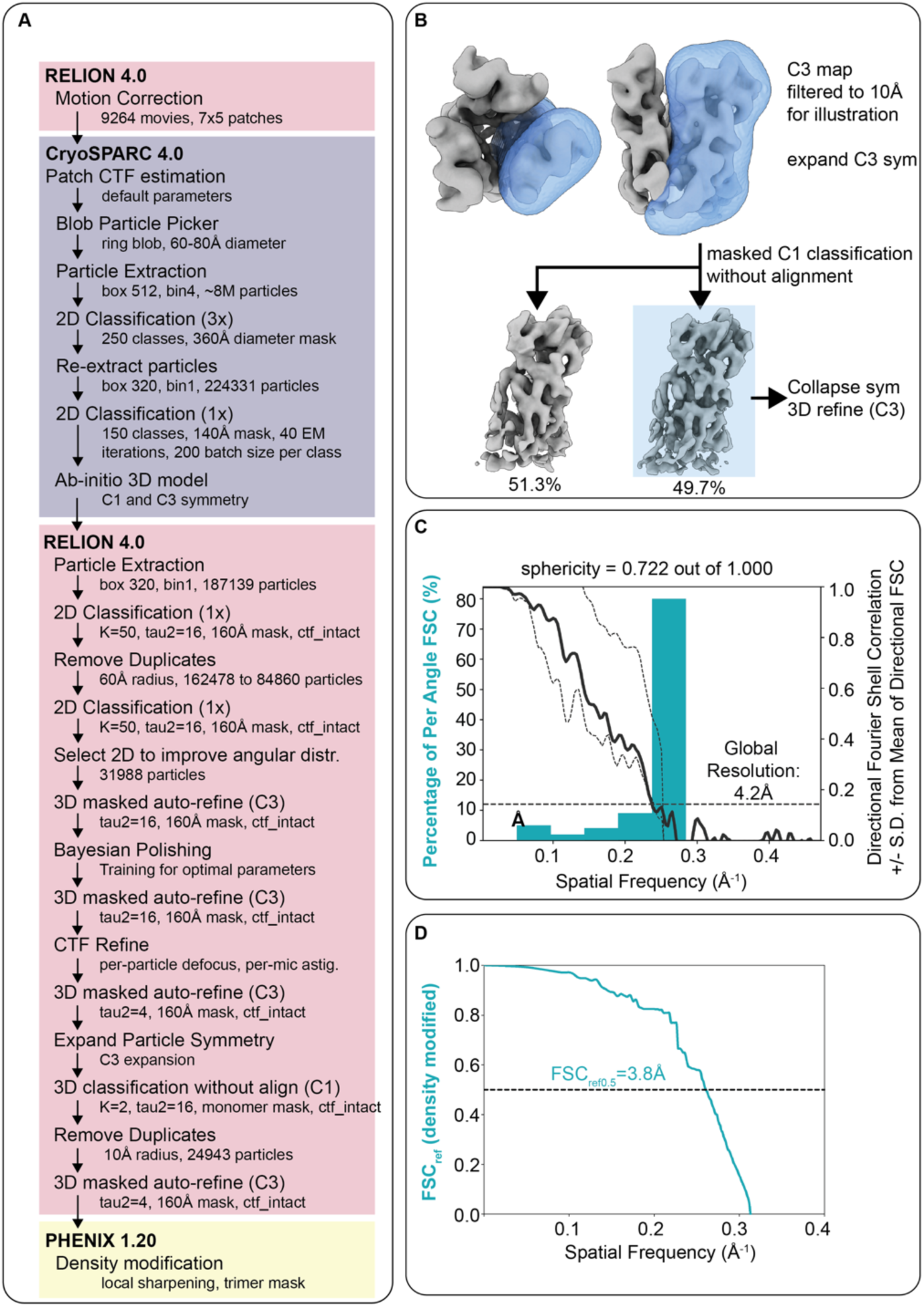
SPA processing workflow of the A10 trimer. **A)** Summary of processing steps used in the SPA workflow. **B)** 3D classification scheme used for the removal of particles containing low-quality asymmetric units within the A10 trimer. The blue mask shown was used for classification, the blue box designates which particles were selected for final refinement. **C)** 3D FSC calculations of the masked RELION (Scheres 2012) half maps. Cyan histogram depicts the fraction of particles that reach the corresponding resolution, and the black curves show global FSC +/− SD of FSCs calculated with extensive angular sampling. Global resolution indicated is at FSC 0.143 cutoff. **D)** FSC_ref_ calculation for the Phenix density modified map with a cutoff value of 0.5 for estimated resolution.

**Figure S4:**
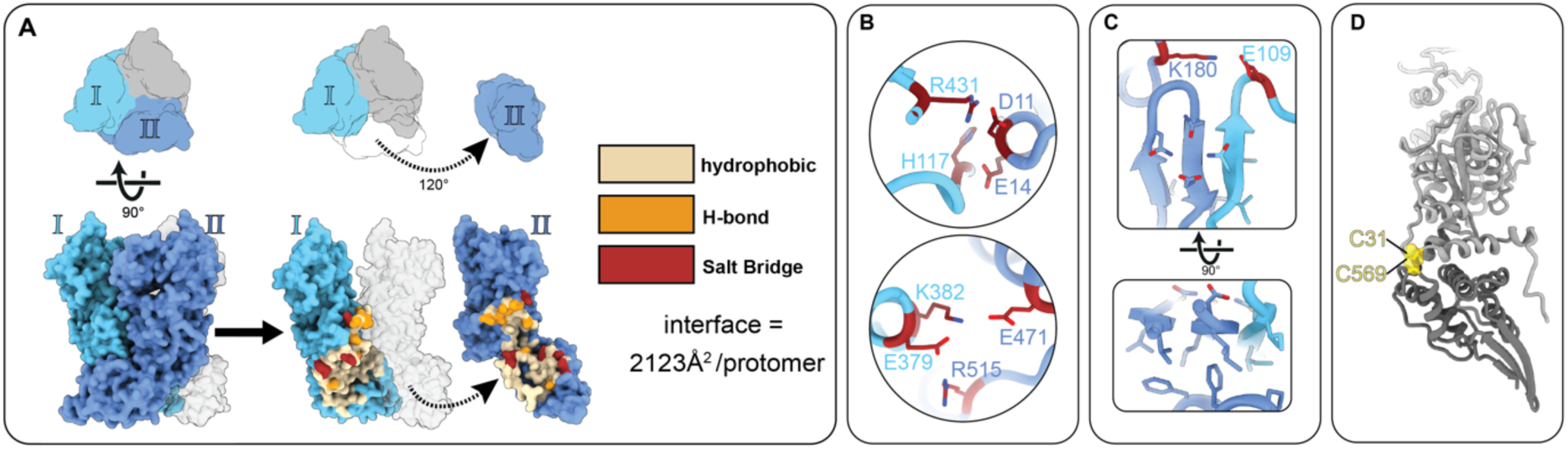
Inter- and intramolecular interactions of the A10 trimer. **A)** Diagrammatic illustration of key inter-chain contacts at the oligomerization interface between two monomers (labeled I and II). The diagram shows monomer II pulling away from the trimer to reveal underlying contacts, shown in yellow (hydrophobic), orange (H-bond), and red (salt bridges). **B)** salt bridge contacts from panel A with residue numbers indicated. Ribbon coloring is the same from panel A. **C)** Top, a salt bridge and hydrogen bond network at the hybrid beta-sheet formed at the core of the trimer. Bottom, hydrophobic packing on the underside of the beta-sheet. **D)** intramolecular disulfide (cysteine residues indicated) which tether the lower C-terminal half of the protein (dark grey) to the upper N-terminal half (light grey). The orientation of the monomer is identical to panel A.

**Figure S5:**
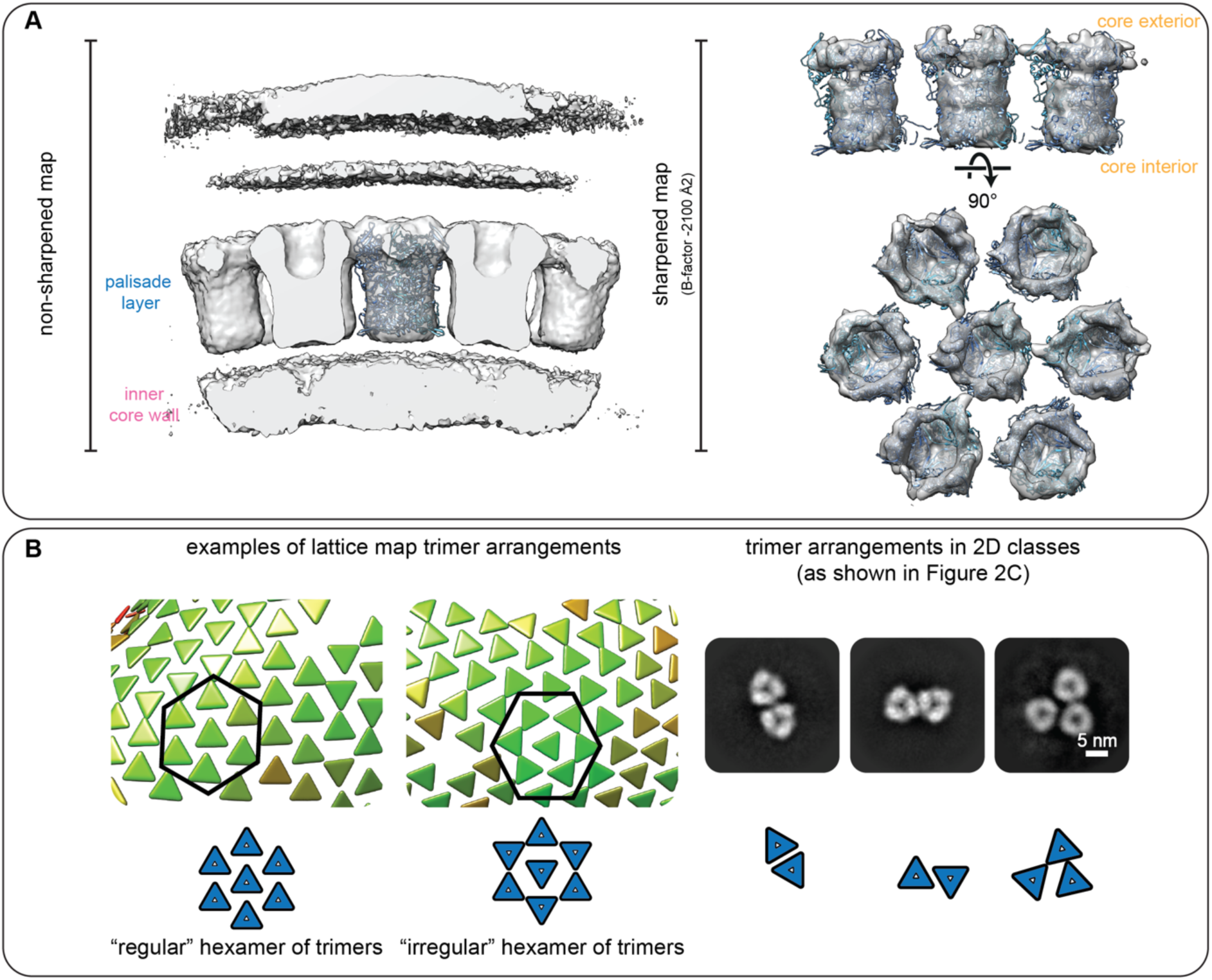
STA results suggest variable trimer interactions within the palisade layer. **A)** Rigid body fit of the A10 trimer into the structure obtained by cryo-electron tomography and subtomogram averaging, both into the unsharpened map (left) and the sharpened map (right). In both cases, it is evident that the A10 trimer is occupying the complete density of the palisade layer. **B)** The variable positioning of the trimers with respect to each other within the palisade layer is shown in lattice maps obtained from STA of cores in intact MV (left) and in 2D classes of trimers released from the core wall of isolated cores (right). The differential orientation of the trimers to each other is always shown schematically. We note that the variable positioning of trimers within the lattice is likely contributing to the limited resolution of our STA average. We also note that the finding of variable trimer interactions in our 2D classes supports that the variable positioning of trimers within the lattice is not due to alignment inaccuracies of STA, but rather indeed suggests the interactions of trimers among each other to be pliable. The color of the triangles denotes the cross-correlation coefficient (CCC) of the alignment ranging from red (low CCC) to green (high CCC). The 2D classes shown here are identical to the ones shown in Figure 2C.

**Figure S6:**
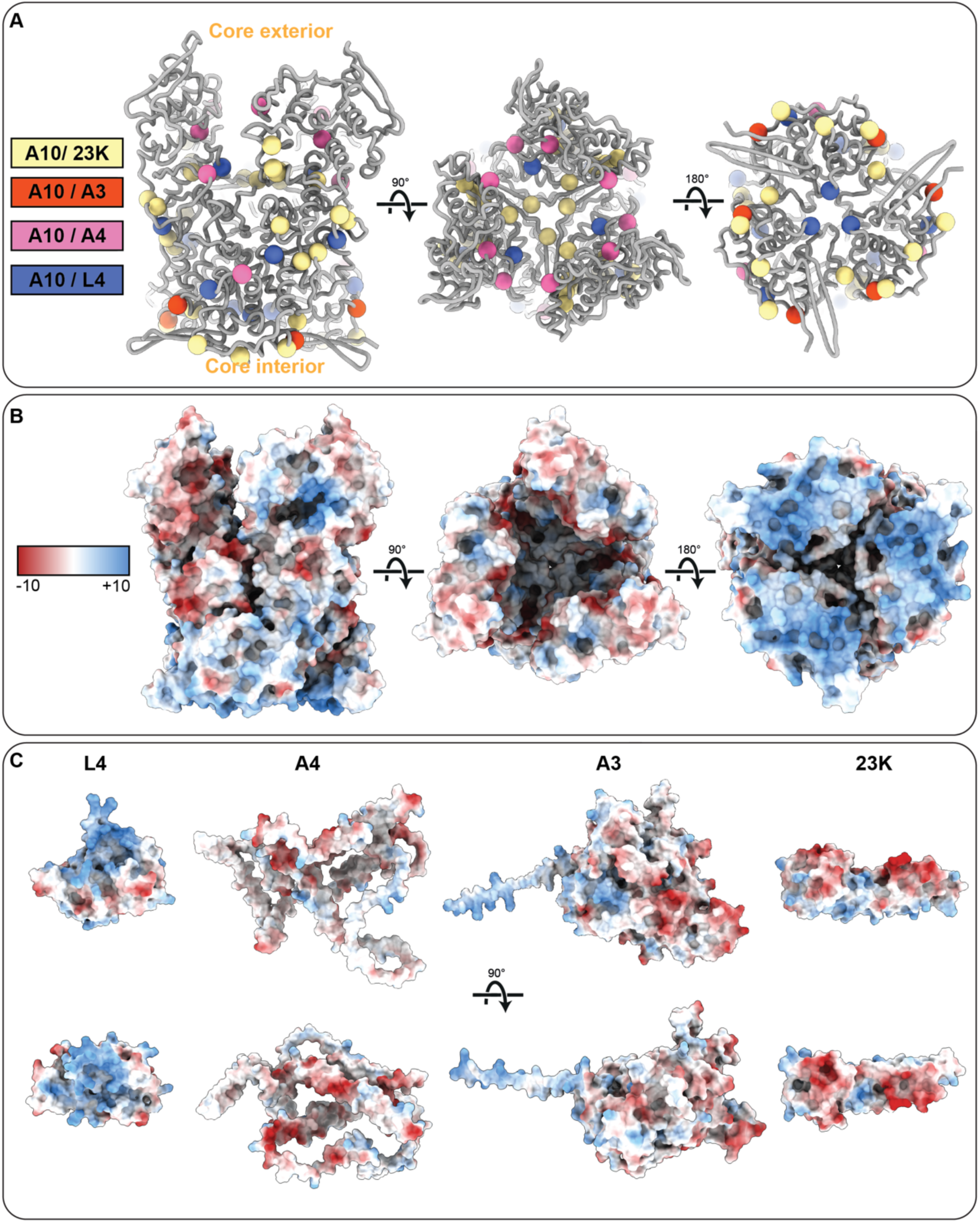
cross-linking mass spectrometry sites on the A10 trimer. **A)** Previously identified (Mirzakhanyan and Gershon 2019) contact sites between A10 and 23K (shown as yellow spheres), A3 (red spheres), A4 (pink spheres), and L4 (blue), as identified via cross-linking mass spectrometry. **B)** Surface charge representation of the A10 trimer (negative charge=red, positive charge=blue). **C)** Surface charge representation of AlphaFold models for monomers of L4, A4, A3 and 23K. Coloring as in (B).

**Figure S7:**
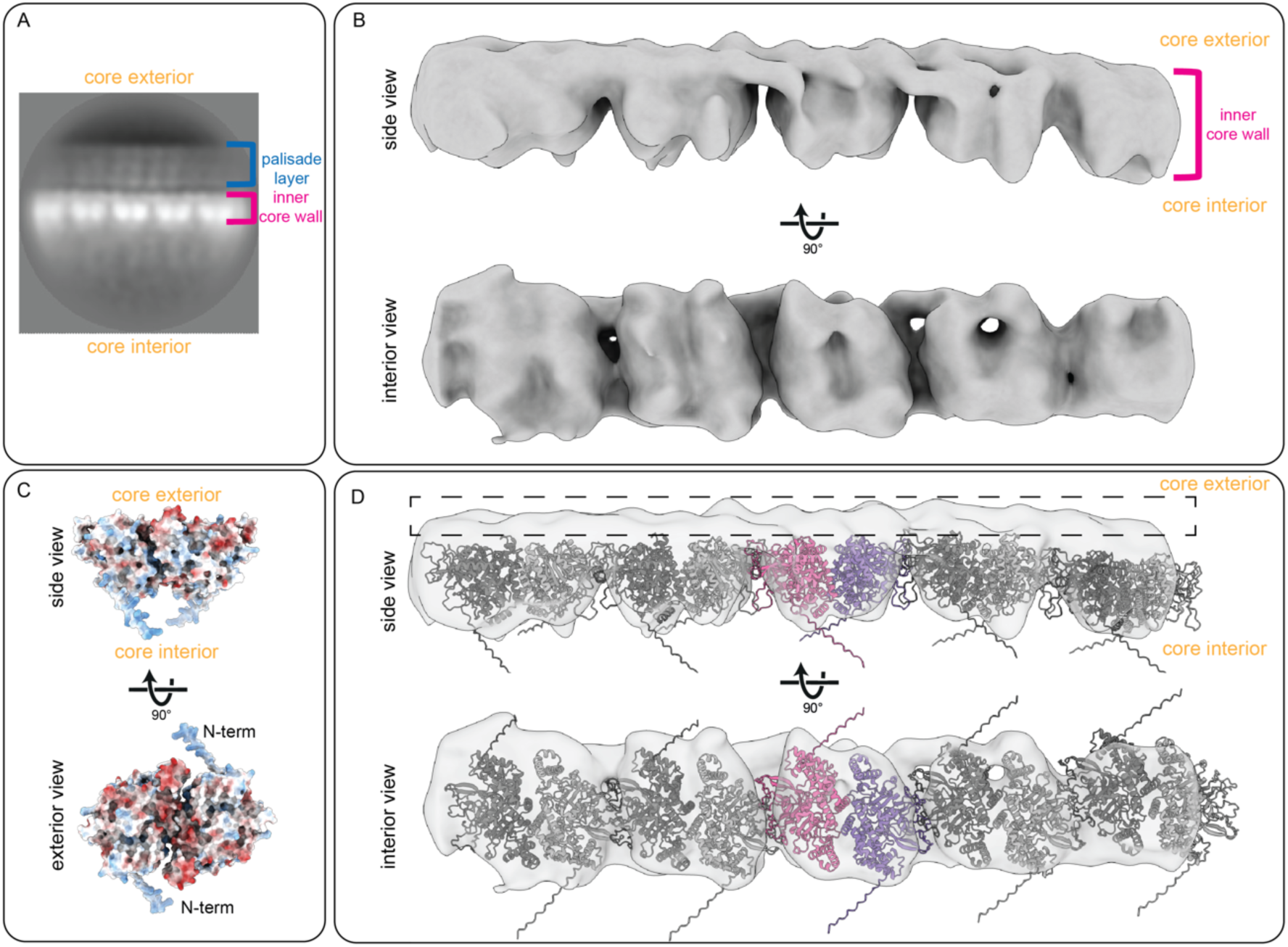
An A3 dimer fits to the density of the inner core wall units. **A)** Projection of the 3D reconstruction of the inner core wall from panel B. **B)** Low-resolution cryo-EM reconstruction of the inner core wall as shown from side and interior views. The variability of the individual units indicates no strict long-range order, as also seen in our 2D classes (Figure 2C) and the top view of the inner core wall in tomograms of isolated cores (Figure 1C). **C)** AlphaFold multimer prediction of an A3 dimer (residues 62-643), with its orientation denoted relative to interior and exterior of virus core. Color represents calculated surface charge with color representations the same as in Figure S6. The orientation was assumed using the negative surface charge of A3 on one side and the suggested orientation towards the base of A10 with the corresponding counter charge (**Figure S6B**). D) Pairs of A3 dimers were rigid body fit into the density of the inner core wall units in panel B. The dimensions and shape of the EM-density accommodate an A3 dimer pair. The central dimer is colored in pink and purple, neighbouring A3 pairs are coloured in grey. The black dashed rectangle annotates a small area of unoccupied density.

**Figure S8:**
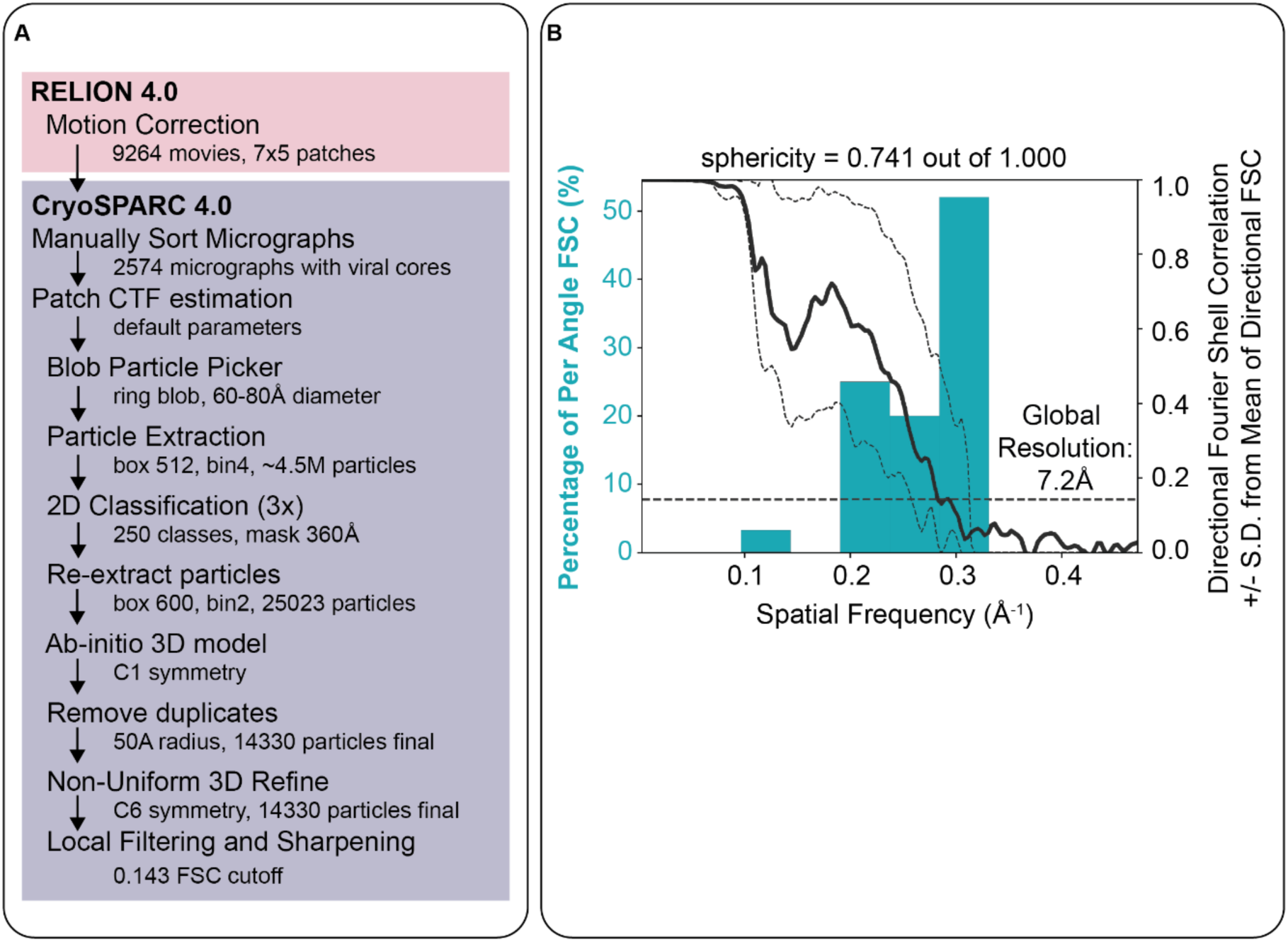
SPA processing workflow of the flower-shaped pore. **A)** Summary of processing steps used in the SPA workflow. **B)** 3D FSC calculations of the masked CryoSPARC half maps. The Cyan histogram depicts the fraction of particles that reach the corresponding resolution, and the black curve shows the global FSC +/− SD of FSCs calculated with extensive angular sampling. The global resolution indicated is at FSC 0.143 cutoff.

**Figure S9:**
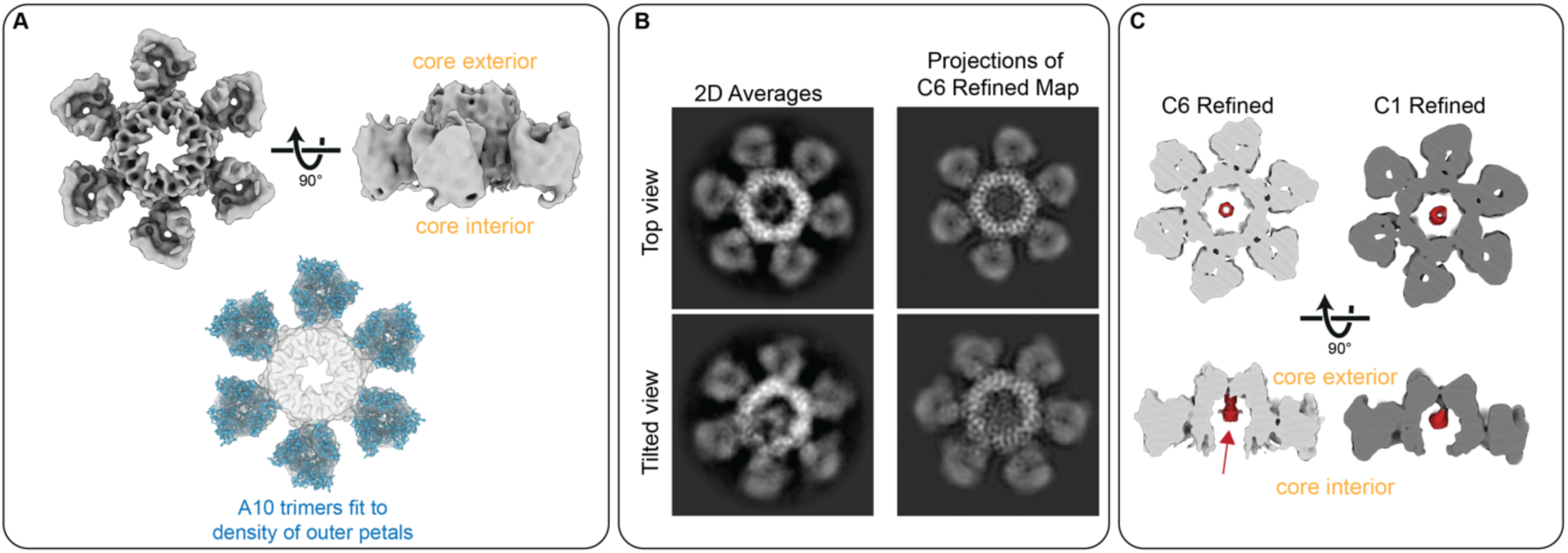
Single particle cryo-EM structure of the flower-shaped pore. **A)** C6-symmetric cryo-EM reconstruction of the flower-shaped pore of the palisade layer. Bottom shows the fit of A10 trimers into the outer densities, while the central hexamer density is not modeled due to the uncertainty of its identity. **B)** Left, Single particle cryo-EM 2D class averages of the flower-shaped pore and right, projections of the C6-symmetrized reconstruction. **C)** Slices through density for C6 and C1 reconstructed flower-shaped pore which reveals an unidentified, donut-shaped central density highlighted in red (red arrow).

**Figure S10:**
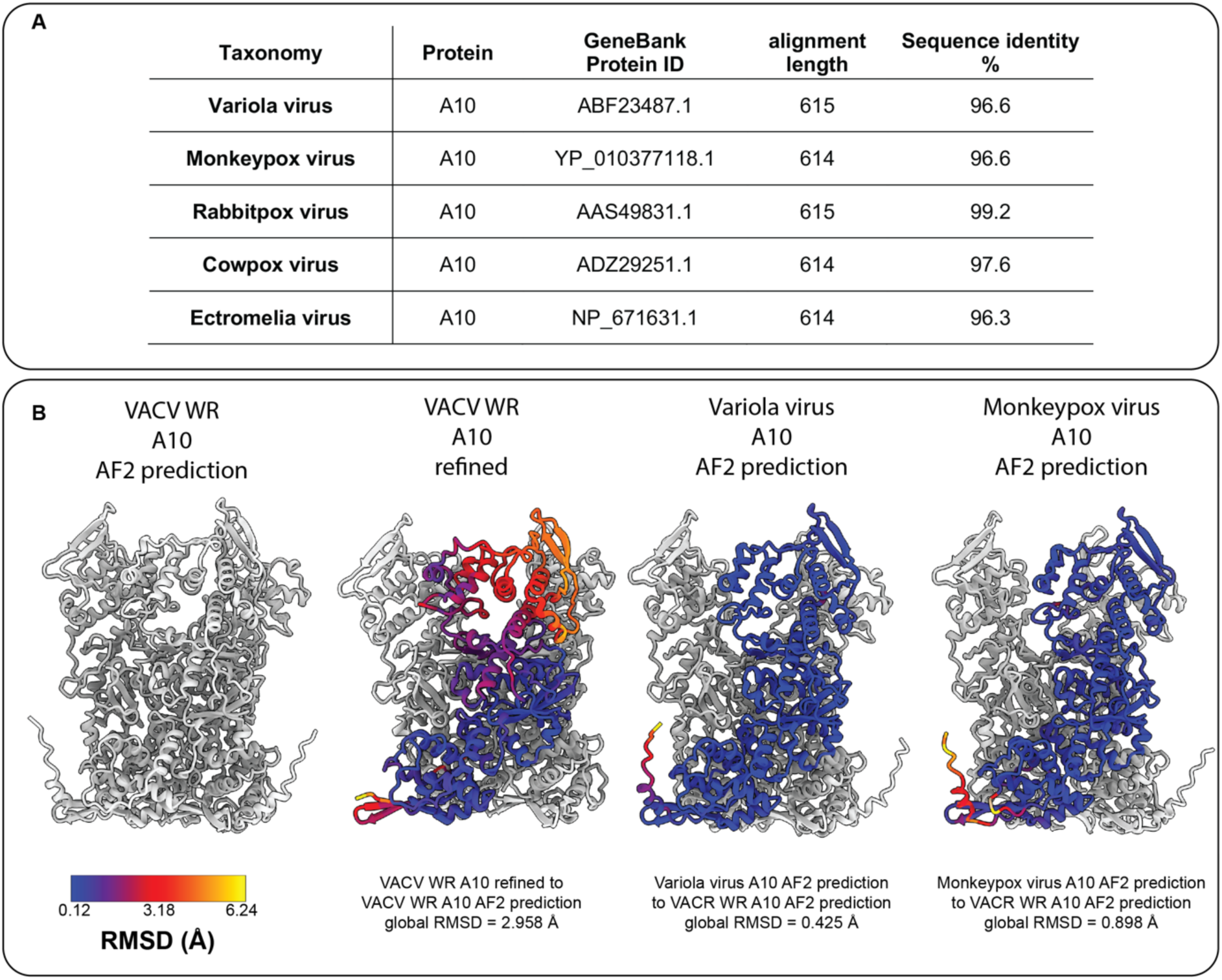
Comparison of Vaccinia virus Western Reserve A10 trimers to other members of the poxvirus family. **A)** Sequence identity of A10 protein of different viruses of the poxvirus family in comparison to VACV WR. **B)** Comparison of the initial VACV WR A10 AlphaFold (AF2) prediction to the refined VACV WR A10 structure (also shown in Figure 3) and AF2 predictions of Variola virus and Monkeypox virus. This comparison shows the strong similarity between the protein folds between Variola virus, Monkeypox virus and VACV WR. It further shows that the biggest difference between the predicted and refined VACV WR A10 model is at the top of the trimer, facing the outside of the core. The color code displays RMSD (root-mean-square deviation) variations, with lower values equaling a more similar structure.

**Figure S11:**
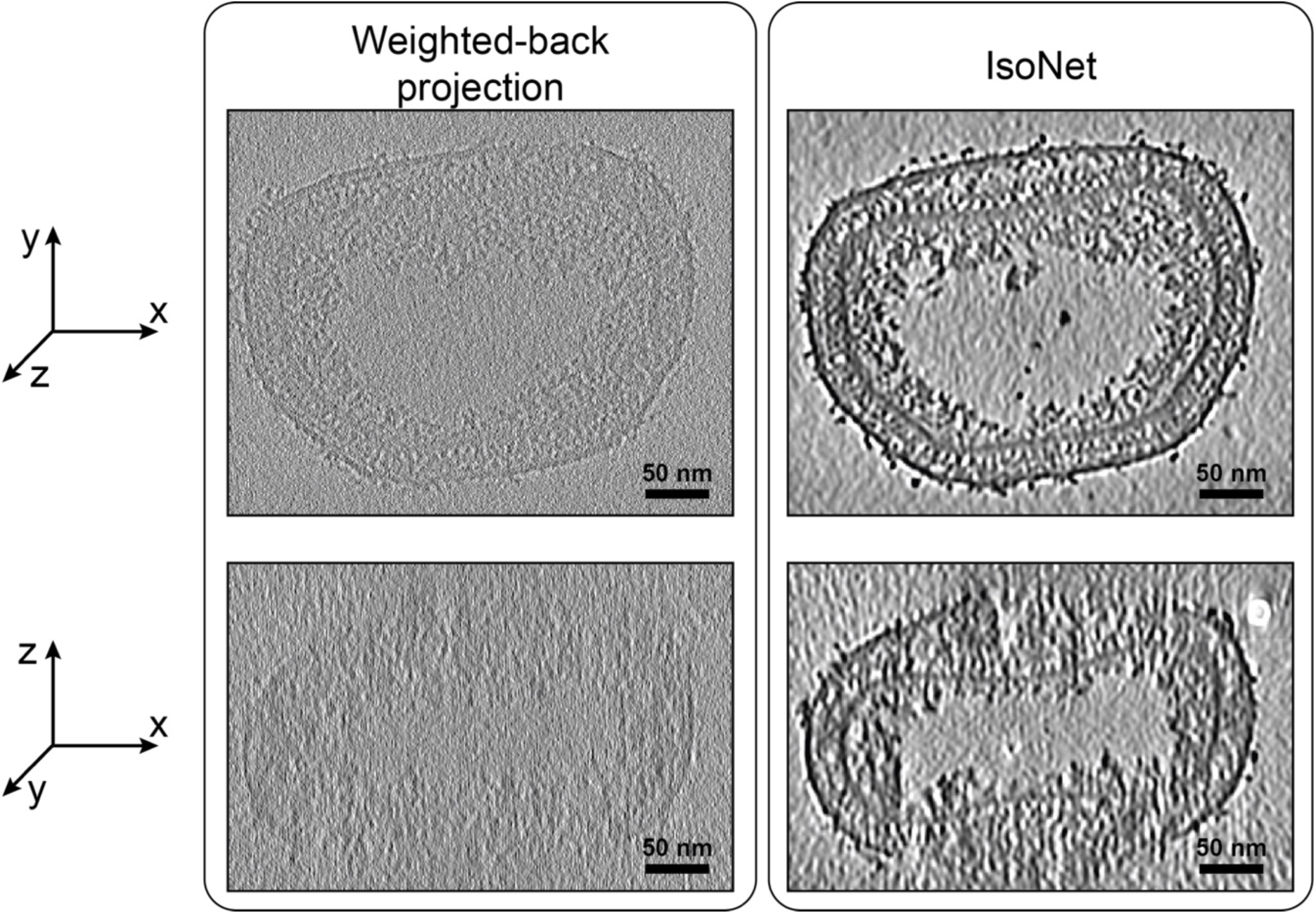
Comparison of weighted-back-projected (WBP) tomogram and IsoNet-corrected tomogram of VACV mature virions. The view axis of the individual slices on the top and bottom of the same tomogram are referenced with a coordinate system on the left, showing the viewing direction onto the xy and xz plane on top and bottom, respectively. The tomogram shown here is identical to the one shown in Figure 1B.

**Table S1:**
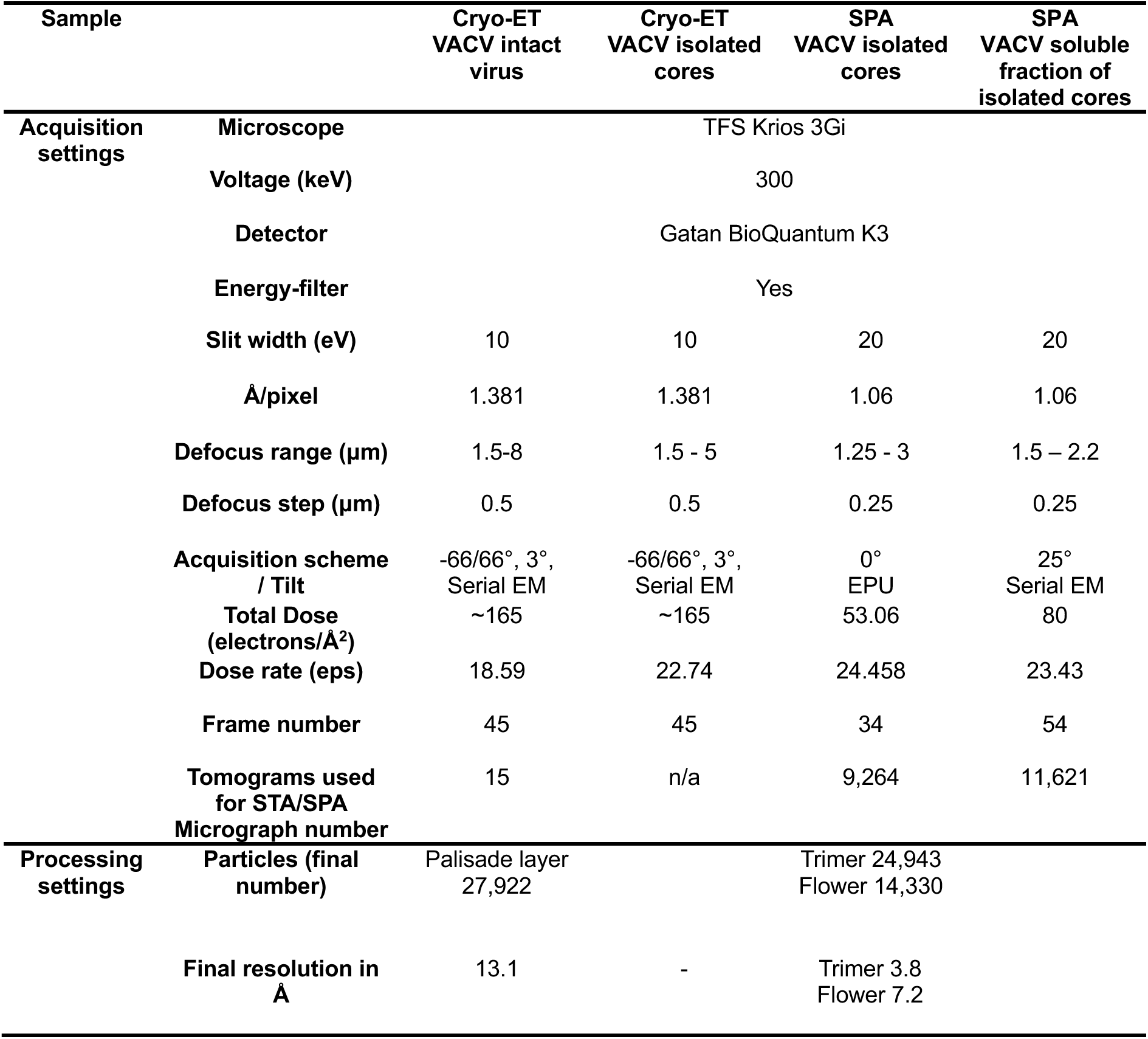
Data acquisition and processing statistics for cryo-ET and SPA.

**Table S2:** Mass spectrometry of components from isolated cores. List of mass spectrometry results from soluble fraction core sample, filtered for VACV proteins. Please note that the precursor proteins are also listed using the UniProt annotation, due to the way of how proteins have been identified computationally within our proteomics data.

| UniProt ID | Common Names | Genes | Sequence Coverage % | Peptide Counts | Log 10 expression |
| --- | --- | --- | --- | --- | --- |
| P16715 | Major core protein A10 (4a) precursor | VACWR129 | 60.5 | 69 | 9.5060 |
| P03295 | Core protein L4 (VP8) | VACWR091 | 54.2 | 16 | 9.1462 |
| P26669 | Cu-Zn superoxide dismutase-like protein A45R | VACWR171 | 91.2 | 9 | 8.8556 |
| P04195 | Cell surface-binding protein | VACWR113 | 53.9 | 21 | 8.7152 |
| P07396 | Phosphoprotein F17 | VACWR056 | 32.7 | 6 | 8.6891 |
| P29191 | A4 (39kDa) core protein | VACWR123 | 17.8 | 4 | 8.2534 |
| P04298 | mRNA-capping enzyme catalytic subunit | VACWR106 | 62.8 | 56 | 8.2447 |
| P68692 | Glutaredoxin-1 | VACWR069 | 24.1 | 4 | 8.2045 |
| P06440 | Major core protein A3 (4b) | VACWR122 | 34.4 | 17 | 8.1866 |
| Q76ZN5 | Profilin | VACWR167 | 39.8 | 6 | 8.0401 |
| P12926 | Core protease I7 | VACWR076 | 24.3 | 11 | 7.8991 |
| P07614 | Protein L3 | VACWR090 | 32.9 | 12 | 7.8896 |
| P04318 | mRNA-capping enzyme regulatory subunit | VACWR117 | 72.1 | 19 | 7.8743 |
| P07242 | Late transcription elongation factor H5 | VACWR103 | 44.8 | 10 | 7.8030 |
| P24758 | Protein A26 | VACWR149 | 35 | 15 | 7.7764 |
| P07616 | Protein J1 | VACWR093 | 19.6 | 4 | 7.7718 |
| P18377 | Phospholipase-D-like protein K4 | VACWR035 | 30.4 | 15 | 7.7231 |
| P68623 | Protein L5 | VACWR092 | 24.2 | 4 | 7.6414 |
| P07611 | Myristoylated protein G9 | VACWR087 | 26.5 | 8 | 7.4532 |
| P68710 | DNA polymerase processivity factor component A20 | VACWR141 | 1.6 | 1 | 7.4508 |

**Table S3:** Model refinement statistics for the A10 trimer.

| **Refinement** | **A10 trimer** |
| --- | --- |
| **Resolution of map (Å)** | 3.8 |
| **Number of A10 monomers** | 3 |
| **Model statistics** |  |
| **All-atom clash score** | 4.96 |
| **Poor rotamers** | 9 (0.54%) |
| **Favored rotamers** | 1644 (98.92%) |
| **Ramachandran outliers** | 6 (0.34%) |
| **Ramachandran favored** | 1755 (97.99%) |
| **Ramachandran distribution Z-score** | -0.02 ± 0.19 |
| **MolProbity score** | 1.26 |
| **Cβ deviations >0.25Å** | 0 (0.0%) |
| **Bonds (RMSD)** |  |
| **Bad bonds %** | 0.00 |
| **Bad angles %** | 0.12 |
| **CaBLAM outliers** | 24 (1.3%) |

**Movie S1:** Video of a cryo-electron tomogram containing an intact purified MV. The shown tomogram corresponds to the tomogram shown in Figure 1B.

**Movie S2:** Video of a cryo-electron tomogram containing an isolated core. The shown tomogram corresponds to the tomogram shown in Figure 1C.

**Movie S3:** Morph between the AlphaFold-predicted A10 trimer, and the refined A10 trimer based on our EM reconstruction

**Movie S4:** Video showing the interactions stabilizing the A10 trimer as described in Figure S4.

